# Continuous transcription initiation guarantees robust repair of transcribed genes and regulatory regions in response to DNA damage

**DOI:** 10.1101/712364

**Authors:** Anastasios Liakos, Dimitris Konstantopoulos, Matthieu D. Lavigne, Maria Fousteri

**Affiliations:** Institute for Fundamental Biomedical Research, BSRC ‘Alexander Fleming’, 34 Fleming st., 16672, Vari, Athens, Greece; Department of Biology, School of Science, National & Kapodistrian University of Athens, Athens, Greece; Department of Biology, University of Crete, 70013, Herakleion, Greece

**Keywords:** Chromatin accessibility, histone modifications, transcription, RNA Polymerase II, DNA damage response, DNA repair, genome integrity, promoter-proximal pausing, pre-initiation complex, PROMPT, eRNA

## Abstract

Inhibition of RNA synthesis caused by DNA damage-impaired RNA polymerase II (Pol II) elongation is found to conceal a local increase in *de novo* transcription, slowly progressing from Transcription Start Sites (TSSs) to gene ends. Although associated with accelerated repair of Pol II-encountered lesions and limited mutagenesis, it is still unclear how this mechanism is maintained during recovery from genotoxic stress. Here we uncover a surprising widespread gain in chromatin accessibility and preservation of the active histone mark H3K27ac after UV-irradiation. We show that the concomitant increase in Pol II release from promoter-proximal pause (PPP) sites of most active genes, PROMoter uPstream Transcripts (PROMPTs) and enhancer RNAs (eRNAs) favors unrestrained initiation, as demonstrated by the synthesis of short nascent RNAs, including TSS-associated RNAs (start-RNAs). In accordance, drug-inhibition of the transition into elongation replenished the post-UV reduced levels of pre-initiating pol II at TSSs. Continuous engagement of new Pol II thus ensures maximal transcription-driven DNA repair of active genes and non-coding regulatory loci. Together, our results reveal an unanticipated layer regulating the UV-triggered transcriptional-response and provide physiologically relevant traction to the emerging concept that transcription initiation rate is determined by pol II pause-release dynamics.

## Introduction

Initiation at transcription start sites (TSSs) of RNA polymerase II (Pol II) and promoter-proximal pause (PPP) release into productive elongation are ubiquitous and crucial steps regulating the transcription of protein-coding genes and long non-coding RNAs^1,2^ (together called mRNAs in this manuscript). The same stands for the transcription of regulatory non-coding regions expressing enhancer RNAs (eRNAs) bidirectionally from enhancer TSSs (eTSSs)^3–5^ and for PROMoter uPstream Transcripts or upstream antisense RNAs (collectively called PROMPTs inhere), which are produced in the opposite direction to mRNA when two stable transcripts are not initiated in very close proximity and in opposite directions^6^. Contrary to mRNAs, eRNAs and PROMPTS are short and their detection can be technically challenging because they are unstable as a result of high early-termination rates and increased susceptibility to degradation by the RNA exosome^6,7^.

Initiation of transcription by Pol II in all the above regions depends on the efficient assembly of the pre-initiation complex (PIC) upstream of transcription start sites (TSSs) and on TFIIH-dependent promoter opening and phosphorylation of serine 5 (S5P) residue in the C-Terminal Domain (CTD) of Pol II^8,9^. After elongation of ∼30–60 nucleotides of initiation-associated RNAs (or TSS-associated RNAs), so-called start-RNAs^10,11^, Pol II is paused at PPP sites by negative elongation factors DSIF and NELF^2,12^. Signal-regulated phosphorylation of these factors and of serine 2 residue (S2P) of Pol II CTD by P-TEFb is required for productive elongation^13–15^. It recently emerged that, if this step does not occur rapidly, start-RNAs are terminated^14,16^, implying that Pol II turnover at PPP sites is high and that replenishment of Pol II engaged in early transcription is achieved by the continuous re-entry of pre-initiating Pol II into PICs^16,17^.

The integrity of the genetic information encoded in DNA sequence is persistently challenged by a variety of genotoxic perturbations^18^. A plethora of DNA Damage Response (DDR) mechanisms have evolved to guarantee the detection and removal of different types of DNA lesions, limiting the probability of mutagenesis by adjusting to the cell’s status and need for efficient recovery from DNA damage^19–21^. Nucleotide Excision Repair (NER) plays a vital role in sensing and removing a large panel of bulky helix-distorting DNA adducts such as Cyclobutane Pyrimidine Dimers (CPDs) induced by ultraviolet (UV) light, as well as benzo[a]pyrene guanine adducts induced by cigarette smoke^19,22^. Transcription Coupled-NER (TC-NER) is promptly triggered by elongating Pol II molecules encountering DNA adducts and speeds-up excision and repair in expressed loci^23,24^. In comparison, the second NER sub-pathway, Global Genome-NER (GG-NER) operates through the entire genome but recognizes more stochastically helix distortions^19,22,25,26^. Importantly, given all the classes of transcripts defined above, it is estimated that the coverage of transcribed regions^27^ potentially scanned by TC-NER expands to more than 50% of the genome, thus qualifying transcription as a major driving force in safeguarding genomic stability.

Although TC-NER depends on lesion-sensing potential by elongating Pol II molecules, transcription elongation has been shown to be transiently inhibited after UV irradiation^28–30^ due to a proportion of Pol II molecules stalling at encountered DNA damages^28,31^. Moreover, depletion of the pre-initiating hypo-phosphorylated Pol II(-hypo) isoform from chromatin shortly after UV irradiation^28,32,33^ has led to the assumption that new transcription initiation events are transiently and globally repressed^32–37^. On the other hand, recent reports^28,29,38^ have revealed a functionally essential stress-dependent global increase in 5’ nascent RNA (nRNA) activity that depends on the UV-induced raise in active P-TEFb levels^39,40^ and on the rapid dissociation of the NELF complex^41^. The ensuing fast and global release of *de novo* Pol II elongation waves from PPP sites into gene bodies boosts lesion-sensing activity and accelerates removal of DNA adducts by TC-NER in virtually all active mRNA genes^28^. Together these findings substantiate the possibility that UV might not as severely affect initiation of transcription, in contrary to previous beliefs.

Taking also into consideration recent evidence that supports the model of disengagement of a given Pol II molecule from DNA template after damage recognition^34,35,42^, it is tempting to assume that ensuring continuity in transcription initiation may bring advantages in the repair process. We thus hypothesized that the apparent loss of pre-initiating RNAPII may not be due to the absence of RNAPII recruitment at TSSs, but rather due to a decrease in the dwell-time of Pol II-hypo isoform at TSS, as justified by the concomitant increase in S5P- and S2P-pol II downstream of TSS^28^. In this way, cells would be able to uninterruptedly feed the global release of lesion-scanning enzymes into transcribable sequences and guarantee the detection of more lesions along DNA template strands.

Herein, we deciphered chromatin dynamics genome-wide upon UV damage and found a significant gain in accessibility (Assay for Transposase-Accessible Chromatin using sequencing, ATAC-seq) at the TSSs of virtually all active regulatory regions controlling mRNAs, PROMPTs and eRNAs expression. This phenomenon was underlined by the maintenance of active histone marks (H3K27ac), the lack of deposition of transcriptional silencing modifications (H3K27me3) and correlated with the influx of Pol II into productive elongation. The paradoxical decrease in pre-initiating Pol II-hypo at these TSSs upon UV was elucidated by revealing that the presence of Pol II-hypo could be rescued when PPP release was drug-inhibited. Accordingly, preserved production of start-RNAs after UV stress lied under the increased production of nRNA and was prevented only after inhibition of transcription initiation. The identified genome-wide dependence of initiation rate on promoter-proximal pause release dynamics explains the seamless recruitment/initiation of Pol II upon UV, in turn enabling efficient repair of the totality of the sequences encoding active regulatory regions and mRNAs.

## Results

### Chromatin accessibility increases at active regulatory regions upon UV irradiation

To characterise the impact that UV might have on the chromatin landscape of transcriptional regulatory regions and how this could be linked to the widespread PPP-release of elongating Pol II and the local increase in nRNA production downstream of TSS^28,29,38^, we first determined the genome-wide changes in chromatin accessibility. The omni-ATAC-seq protocol^43^ was implemented in our system involving the irradiation with mild doses of UV-C of human skin fibroblasts synchronized in early G1 (see Methods and also^28^). We reproducibly measured chromatin accessibility before (NO UV) and after (+UV) irradiation during the early phase of recovery (Supplementary Fig. 1a) and mapped a total of 105,574 Accessible Regions (ARs) across conditions. ARs were enriched at promoters and intragenic or intergenic regions with transcriptional regulatory function (TSSs, TSSs flanks and enhancers according to ChromHMM annotation, Fig. 1a, Fig. 1b,c, Supplementary Fig. 1 b-d, Methods). Interestingly, we reveal a widespread increase (up to 1.71 average Fold Change (FC)) in chromatin accessibility after stress at 97.9% of promoter-, 94.6% of intragenic- and 94.4% of intergenic-ARs (Fig. 1 b-d, Supplementary Fig. 1 d, e).

**Fig. 1.**
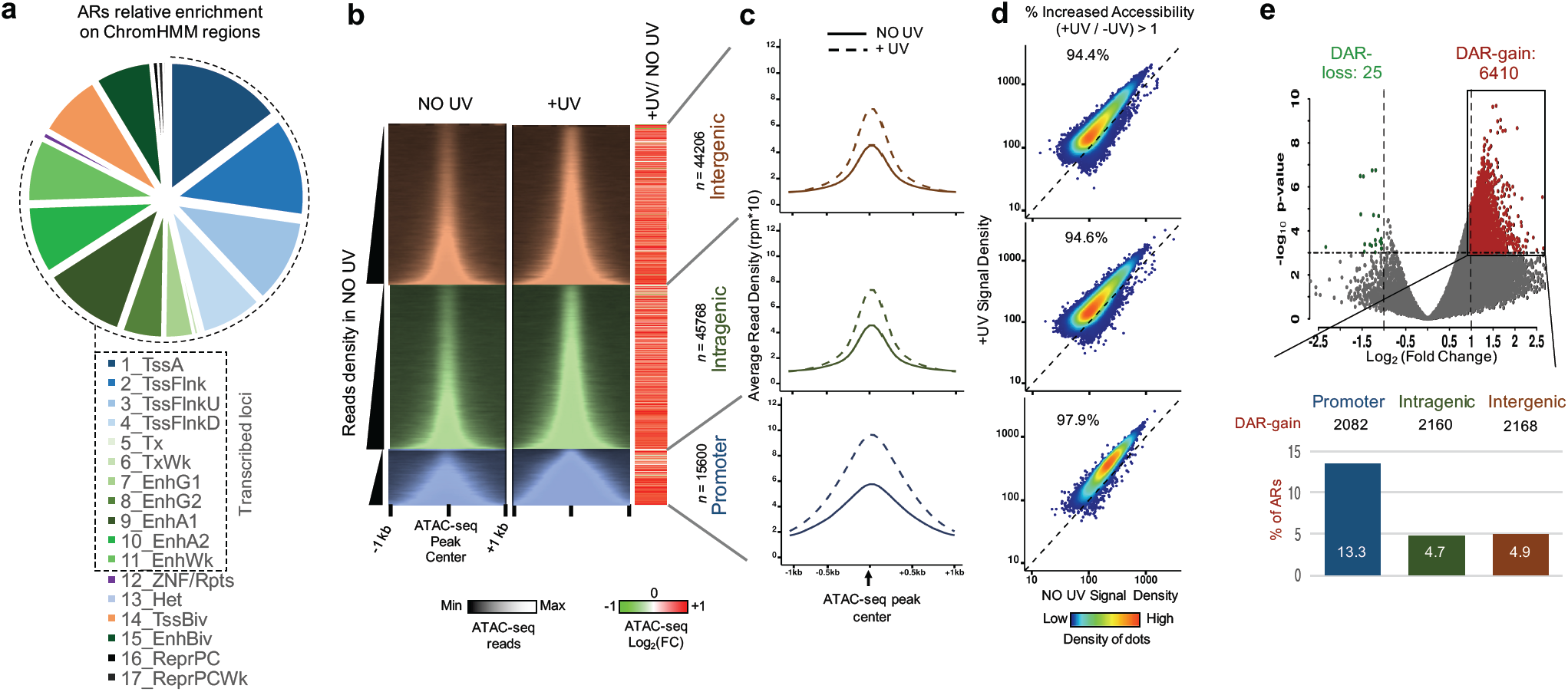
Increase of chromatin accessibility in response to mild doses of UV irradiation. **a** Classification of ARs according to ChromHMM annotation. Dashed line represents active regulatory loci. **b** (Left panel) Heatmap of ATAC-seq reads in genomic regions 1kb around ATAC-seq peak centres before (NO UV) and after UV (+UV), categorised according to their genomic position relative to RefSeq genes (intergenic, intragenic and promoter peaks) and sorted by increasing read density (as determined before UV). (Right panel) Heatmap showing the log_2_ Fold Change (log_2_ FC) between +UV and UV read densities calculated in genomic regions 1kb around ATAC-seq peak centers. **c** Average profile plots of ATAC-seq read densities of non-irradiated (solid line) and irradiated (dashed line) cells in intergenic (red), intragenic (green) and promoter (blue) regions. **d** Heat density scatter plots comparing ATAC-seq read density before and after UV at all accessible regions (ARs) in intergenic, intragenic and promoter regions, respectively. **e** (Upper panel) Volcano plot representing Differentially Accessible Regions (DARs) between irradiated and non-irradiated cells. Regions with significantly increased (DAR-gain) or decreased (DAR-loss) accessibility, are depicted in red and green, respectively. (Bottom panel) Proportion of DAR-gain loci in intergenic, intragenic and promoter ARs

We then selected Differentially Accessible Regions (DARs) by applying stringent thresholds both in terms of FC (Log_2_ FC > 1) and *P*-value (*P* < 0.001) and found that 6410 loci shown particularly increased chromatin accessibility upon UV (DAR-gain) (Fig. 1e, top panel). DAR-gain found at promoter regions represented 13,3% of all promoter ARs (Fig. 1e, lower panel), thus pinpointing towards a potentially functionally relevant chromatin opening at TSS regions. DAR-gain located at intragenic and intergenic loci (Fig. 1e) were linked to genes if they overlapped functional enhancers defined in FANTOM5 (Methods). We found that genes associated with DAR-gain loci (either identified on their promoter or enhancers) were representative (adjusted (adj.) *P* < 0.05) of a number of biological pathways previously associated with DDR processes, including cellular response to stress, DNA repair, transcription regulation by TP53 and cell cycle checkpoints (Supplementary Fig. 2). In addition, we identified a broad range of many other significant GO categories (163 in total, Supplementary Fig. 2), a result in line with the previously reported global PPP release of elongating Pol II waves at all active gene bodies upon UV irradiation^28^.

### Chromatin marks associated with transcription status remain stable after damage

A number of studies have demonstrated that the turnover, modification and/or degradation of histones around damage sites represent essential steps in conserved pathways that help cells deal with genotoxic stress^44–46^. However, especially in the case of UV-C induced DNA damage, little is known about the post-translational modifications (PTMs) of histones around transcriptional regulatory regions. To better interpret the increase in chromatin accessibility and clarify its possible impact on genome-wide transcription dynamics, we studied the differential presence of two histone PTMs representative for the transcription status of associated chromatin: the silencing mark H3K27me3 and the activation mark H3K27ac^47–49^.

We conducted ChIP-seq experiments with antibodies specific for these histone PTMs in NO UV and + UV conditions and focused our analysis on TSSs of mRNAs and with on a robust set of eTSSs, which are known to be functional and potentially transcribed in the investigated cell type according to the FANTOM5 database. We used the ChIP-seq data generated in this study (H3K27ac and H3K27me3), as well as previously published ChIP-seq data (Pol II-ser2P^28^) from the steady-state (NO UV) condition, to determine subsets of Active (presence of H3K27ac and Pol II-ser2P peaks over TSS), Repressed (presence of H3K27me3 peaks over TSS) and Inactive loci (no peak detected over TSS for H3K27ac, H3K27me3, and Pol II) (Fig. 2a, see Methods) in our cell system. We associated the changes in histones marks and Pol II observed in these regions upon UV with ATAC-seq results. The increase in chromatin accessibility was detected at all active TSSs, which corresponded largely to the promoters identified above (compare Fig. 1a and Fig. 2a, and see Methods), as well as FANTOM5-annotated active eTSSs upon UV (Fig. 2b, ATAC, 95% Confidence Interval (CI) excludes 0). This opening was in sharp contrast to the UV-induced global loss of Pol II-hypo at TSSs and eTSSs (Fig. 2b, Pol II-hypo, 95% CI excludes 0) (Fig. 2a and b, and Supplementary Fig. 3) at these regulatory regions.

**Fig. 2.**
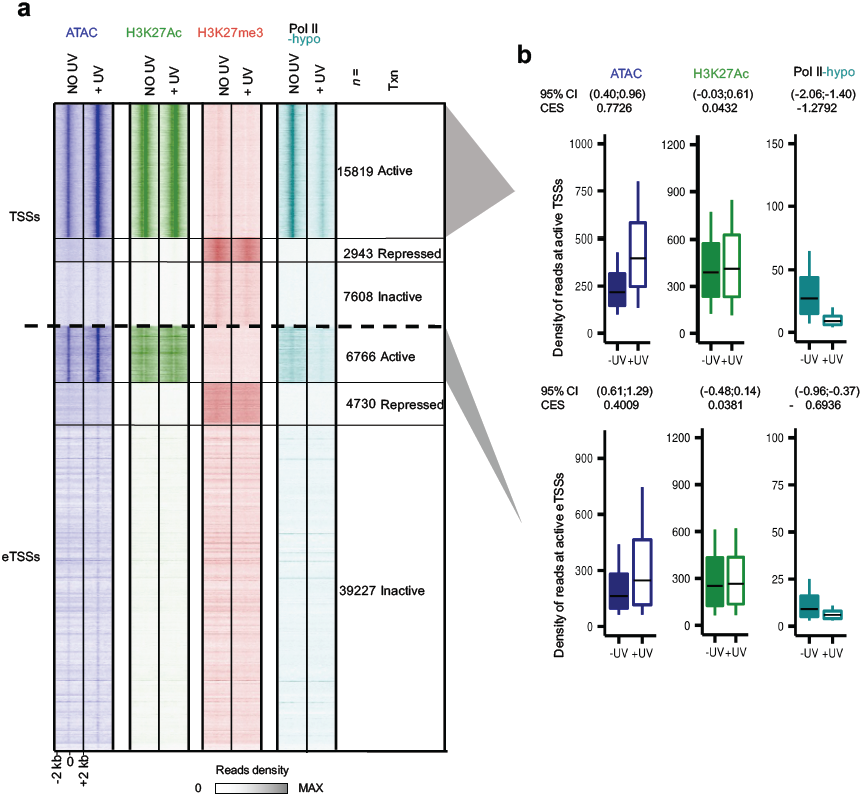
Chromatin modifications remain virtually stable upon UV damage. **a** Heatmap depicting read densities for ATAC-seq, H3K27ac, H3K27me3 and Pol II-hypo ChIP-seq before (NO UV) and after UV (+UV), for genomic regions 2kb around active, inactive and repressed TSSs and eTSSs, respectively (see Methods). Data for Pol II-hypo are obtained from^28^ **b** Box plots summarizing quantifications of ChIP-seq reads shown in **a** for active TSSs and eTSSs, respectively. Box plots show the 25th–75th percentiles, and error bars depict data range to the larger/ smaller value no further than 1.5 * IQR (inter-quartile range, or distance between the first and third quartiles). 95 % confidence intervals (CI) of mean differences between + UV and NO UV of log_2_ counts were calculated for 10,000 samplings of 100 data points with replacement from each population. Effect sizes of log_2_ counts between irradiated and non-irradiated samples were calculated using Cohen’s method (CES)

Strikingly, we also found a high stability in the levels of H3K27ac (Fig. 2a, b, 95% CI includes 0, and Supplementary Fig. 3) and we observed no exchange of H3K27ac for H3K27me3 in response to UV at these active TSSs and eTSSs. Reciprocally, there was no loss of H3K27me3 for H3K27ac, and no gain of Pol II at repressed loci (Fig. 2a, Supplementary Fig. 3). Accordingly, the results of our genome-wide analysis were consistent with biochemical evidence obtained by histone acetic extraction followed by Western Blot analysis, showing that the global levels of H3K27me3 or H3K27ac remain fairly stable during the full period of recovery from UV stress (Supplementary Fig. 3c, d).

We therefore conclude that depletion of detectable Pol II-hypo at TSSs and eTSSs does not occur due to repression of these loci by tri-methylation of H3K27^50,51^, or loss of activating histone mark H3K27ac^48^.

### Chromatin opening parallels Pol II transition into elongation upon UV irradiation

To elucidate the functional advantage associated with increased chromatin accessibility in response to UV, we performed a thorough integrative analysis of our data in relation with previously published datasets (Pol II-ser2P from^28^and CAGE-seq from^4^, see Methods). First, we customised a genome annotation, which unambiguously pinpoints to the TSSs of mRNAs, PROMPTs, and eRNAs that do not overlap with regions possibly being transcribed through from neighboring/overlapping genes, promoters or enhancers (see Methods). We then established three categories (Fig. 3a-c), as per previously suggested models^52^: first, active bidirectional promoter regions, which include the TSSs of mRNA-mRNA pairs transcribed in opposite directions (Fig. 3a); second, active unidirectional promoters, which include the TSS of one mRNA gene (+ or -) for which we could associate an expressed PROMPT in the antisense direction (Fig. 3b); third, active intergenic—as opposed to intragenic—enhancers to avoid potential contamination by interfering reads that derive from overlapping transcription of other active elements (Fig. 3c). Importantly, PROMPT and enhancer transcriptional activity was defined from available Cap Analysis Gene Expression (CAGE) data for skin and dermal fibroblasts (FANTOM5 consortium, see Methods) that accurately determine transcript starting position (5’ end), abundance and directionality of Pol II transcription in our model (Fig. 3a-c, CAGE). TSS loci were sorted by interCAGE distance, which we defined as the distance separating the summits of CAGE signals detected on the (+) and (-) strands (Fig. 3a, b and Methods). This allowed us to identify regions with overlapping (convergent, CONV) or non-overlapping (divergent, DIV) transcription (Fig. 3a, b). By focusing on the latter category, we could study the dynamics of transcription at play in each direction, without having to deal with potential interferences.

**Fig. 3.**
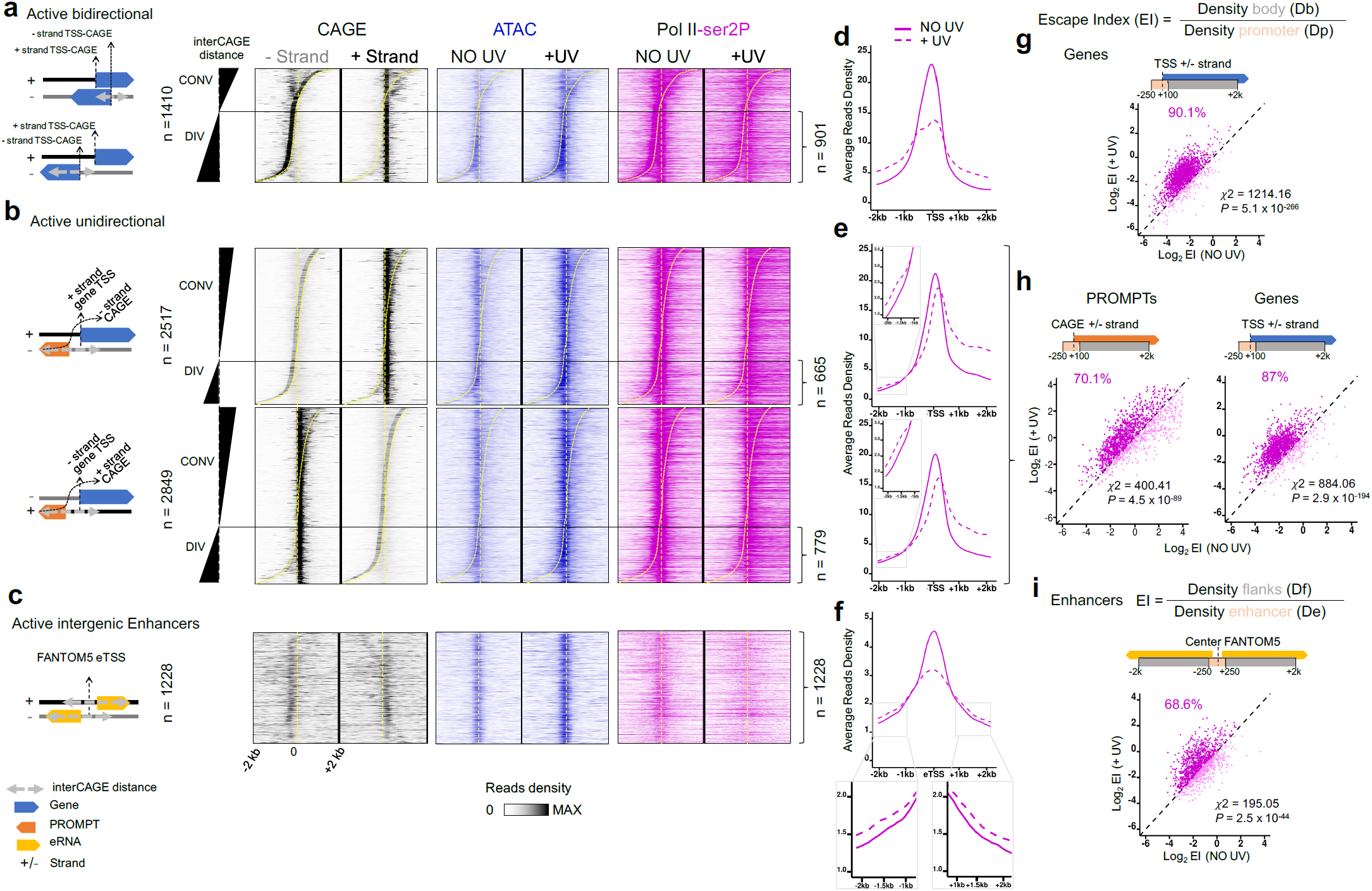
Release of Pol II from pausing sites in coding and non-coding transcribed regions upon UV stress. **a** (Left) Scheme of convergent (“CONV”, overlapping) and divergent (“DIV”, non-overlapping) active bidirectional promoters expressing two mRNAs (Blue arrows). (Right) Heatmap showing distribution of CAGE (black; – and + strand separately) in steady state, and ATAC-seq (blue) and Pol II-ser2P (purple) read densities, before (NO UV) and after UV (+UV) in genomic regions 2kb around + strand TSS-CAGE position. Genomic loci are sorted by interCAGE distance = “+ strand TSS-CAGE” – “-strand TSS-CAGE” (see Methods for details). Divergent loci correspond to interCAGE > 100 bp. **b** Same as in **a** but for active unidirectional promoter TSSs, where PROMPTs (orange arrow) are transcribed in the opposite direction to mRNA gene, from either the – strand (Upper panel) or the + strand (Bottom Panel). Straight dashed lines indicate the position of mRNA TSS, and the sigmoidal dashed line indicate the variable relative position of CAGE-PROMPT. Genomic loci are sorted by interCAGE distance = “TSS-mRNA” – “CAGE PROMPT” and categorised in convergent (mRNA and PROMPT overlapping, interCAGE < 100 bp) and divergent (mRNA and PROMPT non-overlapping, interCAGE > 100 bp) loci. **c** (Left panel) Scheme of active intergenic enhancers expressing eRNAs (yellow arrow) in opposite direction. (Right panel) same as in **a. d**,**e** Average read density profile plots of Pol II-Ser2P before (solid line) and after UV (dashed line) on divergent categories defined in **a**,**b**. Insets represent a zoomed view of the light grey boxes. **f** Same as in **d**,**e**, but for all active intergenic enhancers. **g-i** Comparison of Escape Index (EI, as defined in the Figure), before and after UV for indicated categories of **d, e, f** (see Methods). Percentages of loci with increased EI after UV (dark purple) are shown. Chi-square test between active bidirectional (*n* = 1806) and inactive unidirectional genes (*n* = 5641) **(g)**, active unidirectional genes (*n* = 1444) and inactive unidirectional genes (n=5,641), active PROMPTs (*n* = 1444) and inactive PROMPTs (*n* = 5641) **(h)**, and between active enhancers (*n* = 1228) and inactive enhancers (n=23,026) (i) were performed to determine whether observed number of genes with ΔEI > 1 (number of genes with EI after irradiation greater than EI before irradiation) differs from expected value purely by chance

Using this set-up, we discovered that the UV-dependent increase in chromatin accessibility (Fig. 3 a-c, ATAC) was paralleled by the transition of Pol II into active elongation (Fig. 3 a-c, pol II-ser2P), not only at flanking mRNAs (Fig. 3d, e), but also at adjacent PROMPTs and eRNA sequences (Fig. 3e, f), as shown by the loss in Pol II reads at TSSs and the gain of reads in downstream regions. These results were confirmed quantitatively by showing that Escape Index (EI) of elongating pol II (inverse of Pausing Index^11^) increased in the +UV condition in comparison to NO UV for 90.1% of bidirectional promoters (Fig. 3g, Chi-square test *P* = 5.1 × 10^−266^), as well as for 70.1 % of PROMPTs (Fig. 3h, Chi-square test *P* = 4.5 × 10^−89^) and 68.6 % of eRNAs (Fig. 3i, Chi-square test *P* = 2.5 × 10^−44^). We conclude that the PPP release of Pol II upon genotoxic stress is synchronously triggered at all active transcription units and coincides with increased chromatin breathing. These data extend the previously characterised transcription-driven genome surveillance mechanism^28^ to essential all active gene regulatory regions and give mechanistic insights into the synergy between the increase in chromatin accessibility and the transcriptional response observed upon UV.

### DRB rescues the post-UV detection of Pol II in PIC

We noted that although 63.65 % of the transcribed genome shows reduction in transcription activity (coverage of the transcriptome with Log_2_ FC (+UV/NOUV)) < 0, see Methods), a local increase in nRNA synthesis downstream of TSS of all active genes is detected during the UV-recovery phase^28–30,38^. This observation combined with the above findings on the UV-induced chromatin opening around virtually all active TSSs, PROMPTs and eTSSs are hardly compatible with the previously suggested model of UV-induced global inhibition of transcription initiation. We thus searched for alternative reasons that could explain reduction of Pol II-hypo levels at active TSSs/eTSSs with increased accessibility after UV.

We performed a set of experiments aiming to determine whether Pol II was actually recruited to TSSs upon UV (proxied by its -hypo isoform). First, as depicted in Figure 4a, we irradiated cells with a mild UV dose and we left them to recover for 2 hours, when the levels of Pol II-hypo have been shown to be severely depleted^28,32,33^. We then applied, or not, an inhibitor of the release of elongating Pol II from PPP (DRB, see Methods). Cells were crosslinked 2 hours after the addition of DRB (or DMSO for the control cells). In accord with the previous reports, in cells that were crosslinked 2 h after UV irradiation in the absence of DRB (+UV / X 2 h), or in cells that were crosslinked 4 h after UV irradiation and had been incubated with DMSO for the last 2 h (+UV / −DRB / X 4 h), we detected only minimal levels of pre-initiating Pol II in total chromatin extracts or at TSSs, PROMPTs, and eTSSs, as revealed by Western Blot analysis (Fig. 4b) and ChIP-seq (Fig. 4c, d), respectively. In contrast, when cells had been incubated with DRB for the last 2 h before being crosslinked at 4 h after UV irradiation (+UV / +DRB / X 4h), we observed a significant rescue of pre-initiating Pol II (-hypo) levels in total chromatin (Fig. 4b, two-sided Student’s t test *P* = 0.0055 compared to “+UV/−DRB/X 4h” and *P* = 0.0156 compared to “+UV/X 2h”). The restoration of pre-initiating Pol II levels was even more pronounced when we focused on the occupancy on active TSSs, PROMPTs and eTSSs, where average read densities detected by Pol II-hypo ChIP-seq after DRB treatment (+UV / +DRB / X 4h) matched the control NO UV levels (NO UV / +DRB / X 4h) (Fig. 4c, d). Therefore, even by blocking the stress triggered transition of Pol II molecules from PPP sites into elongation at two hours post UV, when the prior-to-UV Pol II-hypo levels were almost completely depleted, we were able to reveal the underlying continuous *de novo* recruitment of Pol II-hypo molecules in PICs.

**Fig. 4.**
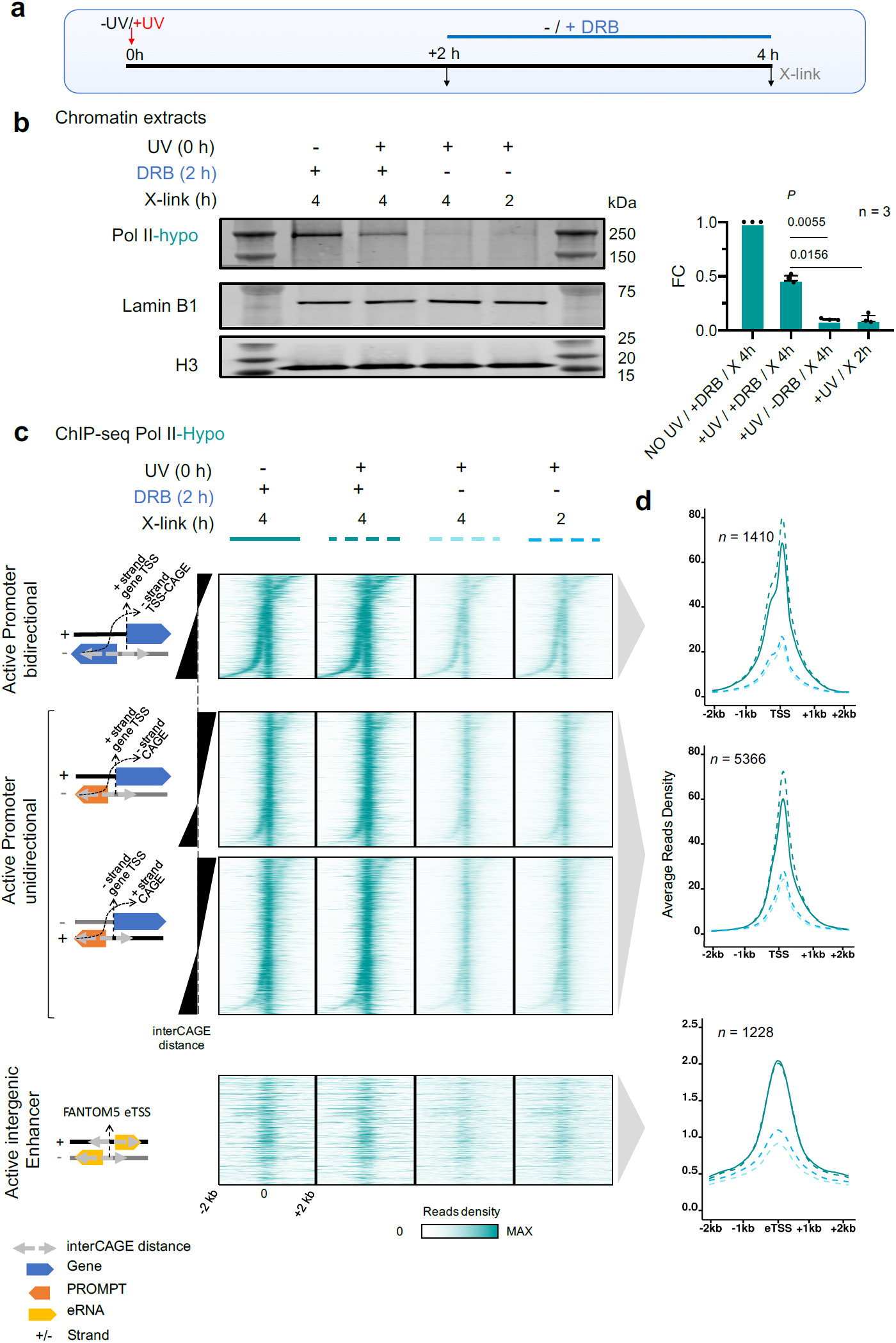
DRB rescues pre-initiating Pol II levels following DNA damage. **a** Experimental timeline showing times of UV irradiation and DRB treatment (see Methods). **b** Western Blot analysis of chromatin extracts for Pol II-hypo levels as examined after employing the experimental strategy described in **a**. Lamin B1 and histone 3 (H3) were used as loading controls. Bar graph represents quantification of Pol II-hypo levels as compared to the NO UV/+DRB condition. Data shown reflect 3 independent experiments. Error bars represent S.E.M. and *P* values are calculated using two-sided Student’s t test. **c** Heatmap of Pol II-hypo ChIP-seq read densities in genomic regions 2kb around TSS for categories defined in **Fig. 3 a-c** after performing the combination of UV/DRB treatments described in **a. d** Average profile plots of read densities analysed in **c**

We also applied DRB just before and for two hours after UV (Supplementary Fig. 4a) and found a limited loss of pre-initiating Pol II in chromatin extracts upon UV (Supplementary Fig. 4b, c, two-sided Student’s t test *P* = 0.0145). This result was corroborated by ChIP-qPCR experiments (performed on the same chromatin extracts used above), as DRB prevented the UV-induced reduction in occupancy of Pol II-hypo at promoter/TSS proximal regions of six active genes (Supplementary Fig. 4d, two-sided Student’s t test *P* = 0.002 for DMSO, while *P* = 0.3138 (non-significant) for DRB).

We thus conclude that the genome-wide UV-induced PPP-release of Pol II molecules into elongation accelerates the transition into initiation of the next-to-be recruited Pol II-hypo molecules, limiting the dwell time of this isoform at essentially all active TSSs, PROMPTs and eTSSs.

### Increased nRNA synthesis from active TSSs upon UV irradiation

Having established that UV irradiation does not inhibit the recruitment of Pol II-hypo into PICs, we next examined the presence of newly synthesized nRNA molecules at TSSs, to determine whether these post-UV recruited Pol II pre-initiating molecules actively proceed into initiation. We took advantage of our and others nRNA-seq data^28,30^ and we examined if the previously characterized global increase of EU- or Bru-labelled RNA reads at the beginning of genes (see Supplementary Fig. 4 in^28^) could originate from increased Pol II initiation at active TSSs (Fig. 5a, b and Supplementary Fig. 5a, b), as suggested before^30^. In particular at unidirectional promoters, we confirmed that nRNA synthesis was increased in the mRNA direction, but we also found a concomitant increase of nRNA production in the antisense, PROMPT direction. Similarly, we found widespread gains in intensity for eRNAs, which emanate equally in both directions from active eTSS (Fig. 5a, b and Supplementary Fig. 5 a, b). Identifying labeled nRNA even at short transcripts such as PROMPTs and eRNAs confirms active labeling close to TSSs and validates the fact that regions directly downstream of TSSs get *de novo* transcribed during the post-UV period. Taken together these data demonstrate that the continuous recruitment of Pol II-hypo molecules (see Fig. 4) and their fast transition into initiation/productive elongation (see Fig. 3), during the post-UV recovery period, is accompanied by synthesis of nascent RNA.

**Fig. 5.**
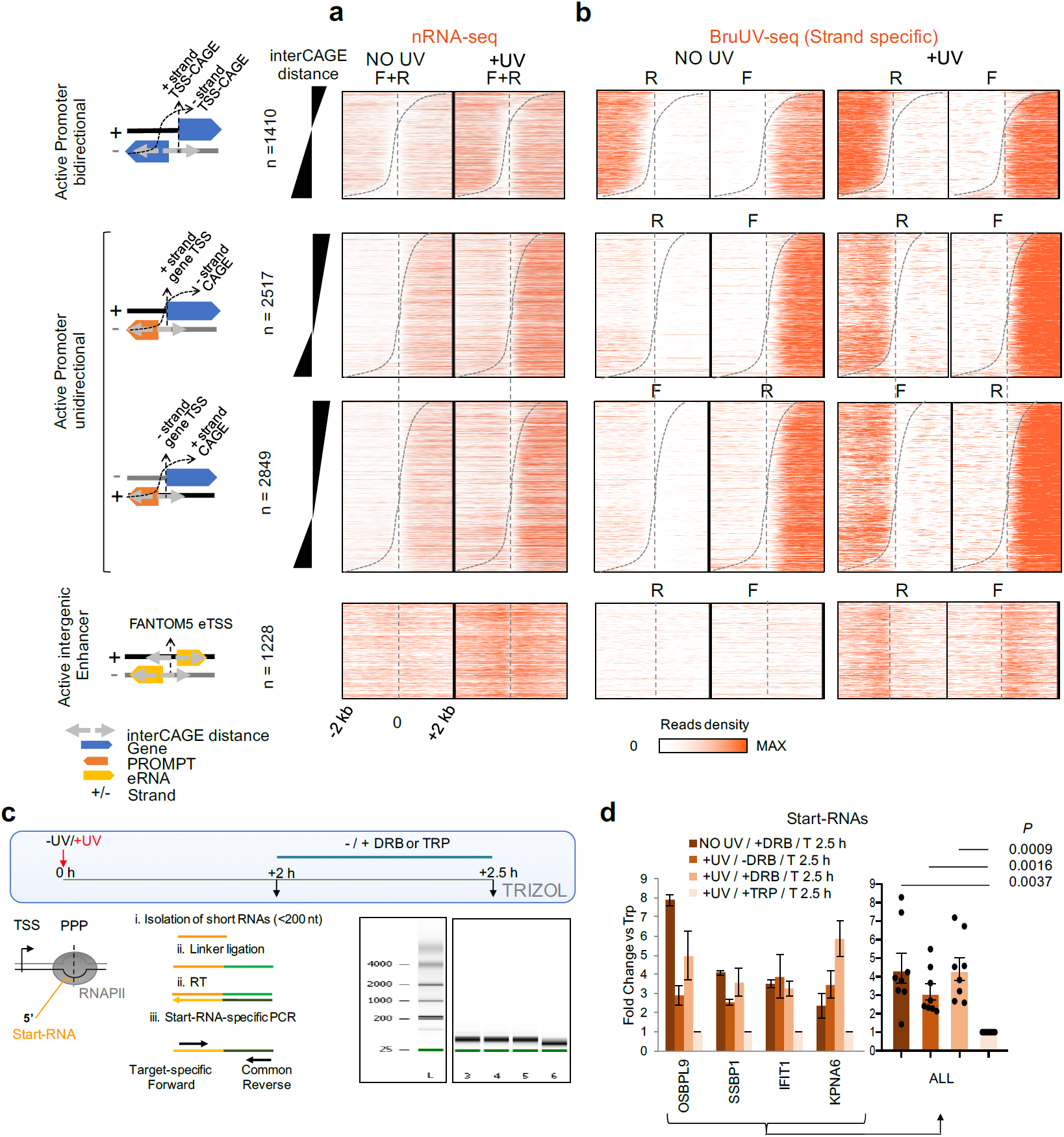
*De novo* and increased RNA synthesis from all TSSs upon UV exposure. **a.** Heatmap of nascent RNA (nRNA) read densities (NO UV and +UV 2 hours data obtained from^28^) before and after UV, in genomic regions 2kb around TSS, for categories defined in **Fig. 3a-c** (see Methods and Supplementary Fig. 5 for time line). F: Forward (+) strand, R: Reverse (-) strand. **b** Same as in **a** for strand-specific BruUV-seq (NO UV and +UV (30 min) data obtained from^30^). **c** (Upper panel) Experimental outline. Cells were treated (or not) with UV and were left to recover normally for 2 h. In turn, DRB, TRP or DMSO was added and after 30 mins cells were disrupted by Trizol addition. (Lower panel, left) Methodology followed for the detection and quantification of gene-specific start-RNAs (for details see Methods). (Lower right) Agilent RNA 6000 Nano Bioanalyzer traces showing size distribution of RNA samples after preparation of a separate small-sized RNA fraction. L: RNA Ladder (size in nucleotides), 3-6: Small-sized RNA fraction of samples analysed in **d. d** qPCR analysis of start-RNAs. Bar chart illustrating FC (compared to TRP treatment) for each gene tested (left) and for the average FC of all genes (right). Error bars represent S.E.M. and *P* values are calculated using two-sided Student’s t test

To further verify initiation activity during UV-recovery, we exploited the possibility to track start-RNAs, which directly inform on the amount of dynamically engaged Pol II located within the initially transcribed sequence (approximately the first 100 nucleotides^11^). We followed the experimental procedure depicted in Fig. 5c and applied, or not, transcription elongation (DRB) or initiation (triptolide-TRP) inhibitors 2 h post UV. For each condition, we isolated small RNAs by size-selection (<200 nucleotides), and we ligated an RNA-DNA linker to their 3’ ends. Reverse Transcription (RT) was performed using a universal primer annealing to the linker sequence as previously described^7^. Subsequently, locus specific qPCR reactions were performed in order to compare, in a quantitative way, the levels of start-RNAs at representative active loci for which we had identified Pol II-ser2P ChIP-seq or nRNA-seq signal (see Methods). Our results revealed that start-RNAs could be detected after UV treatment, validating the fact that initiation still occurs during the UV-recovery phase (Fig. 5d, +UV / −DRB). Similar result was obtained in the presence of the transcription elongation inhibitor (Fig. 5d, +UV / +DRB). However, the opposite was found after inhibiting transcription initiation by TRP, which led to a clear reduction of start-RNAs (Fig. 5d, +UV/+TRP, two-sided Student’s t-test *P* = 0.0037 compared to “NO UV/+DRB”, *P* = 0.0016 compared to “+ UV/−DRB”, *P* = 0.0009 compared to “+ UV/+DRB”), consolidating further evidence of the non-stop recruitment and functional engagement of Pol II at TSSs after UV irradiation.

### Equal levels of Pol II-hypo at PICs primes for uniform TC-NER

Next, we took advantage of XR-seq data (eXcision-Repair sequencing)^24^, which precisely and exclusively pinpoint the location and levels of transcription-dependent repair (TC-NER pathway) when the assay is performed in GG-NER-deficient cells (Xeroderma Pigmentosum (XP)-C cells). Given the strand-specificity of the assay, we considered only the excision of CPD-damages from template (non-coding) strand (TS) for mRNAs, PROMPTs and eRNAs, which corresponded to the + (blue) or the – (red) strand of the genome (Fig. 6a-b) depending on the transcript orientation. Upon correlation with CAGE, we found that onset of TC-NER coincided with CAGE reads location, confirming the fact that TC-NER (triggered by damage-arrested Pol II molecules^24^) and CAGE^4^ accurately locate active TSSs (Fig. 6b (compare with Fig. 3a; left), Fig. 6c, d). As expected, repair efficiency was equal in each direction for bidirectional active promoters (Fig. 6b-d, Fig. 6e, XR-seq (XP-C)). This result was also in line with Pol II-hypo ChIP-seq data showing equivalent amounts of Pol II recruitment at PICs (Supplementary Fig. 6a) and CAGE data indicating balanced production of capped mRNAs (Fig. 6e, CAGE, boxes centered around Log_2_ FC = 0). Nevertheless, we note that the variability between both directions was strikingly less for TC-NER (XR-seq (XP-C)) and Pol II-hypo than for CAGE (Fig. 6e, proportion of non-significant F-Tests: *P* = 0) (Supplementary Fig. 6b, top panel, proportion of non-significant F-Tests: *P* = 0).

**Fig. 6.**
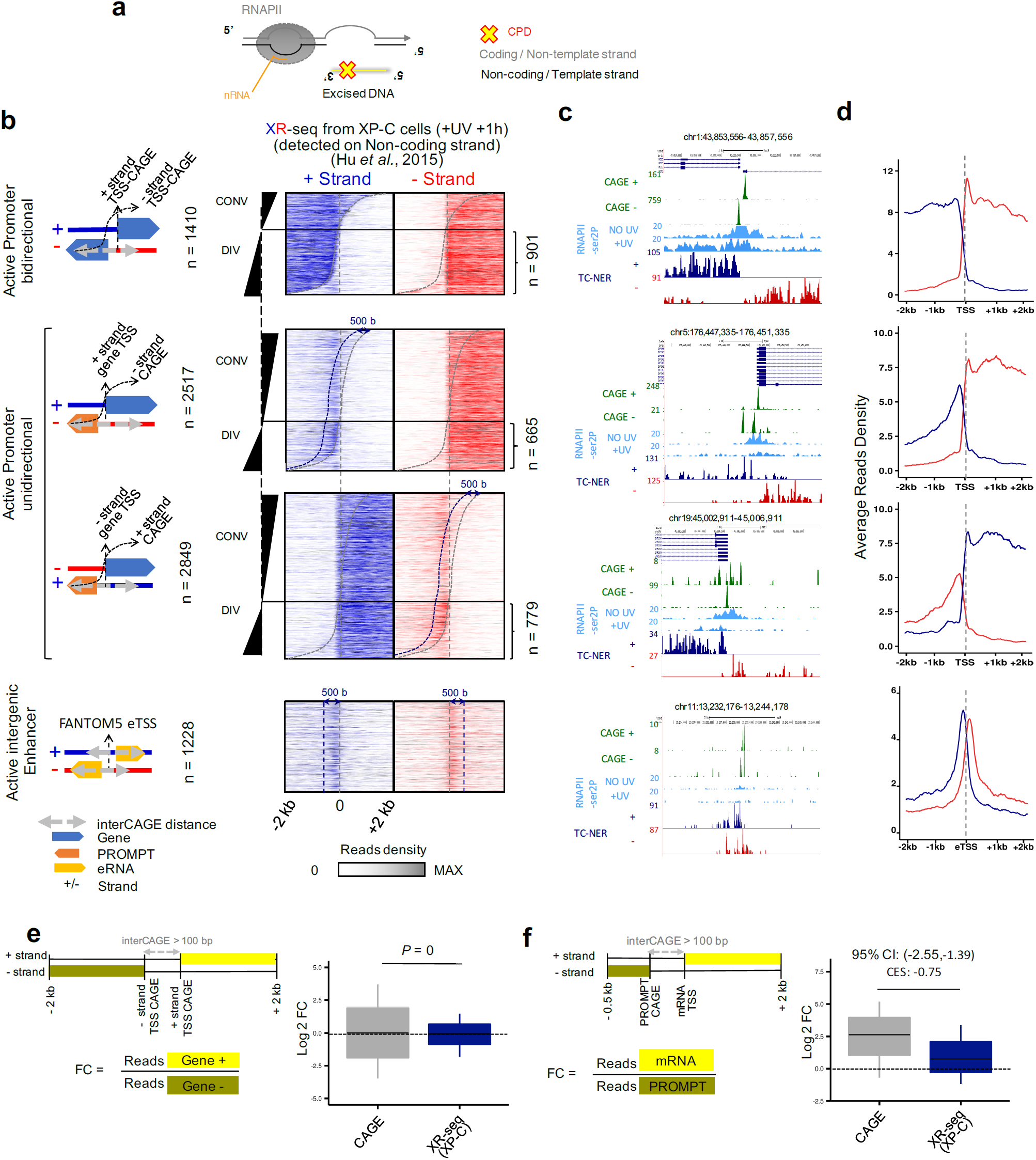
TC-NER is detected homogeneously at all transcribed coding and non-coding regions. **a** Scheme representing the orientation and nomenclature of the DNA strands for TC-NER-specific XR-seq analysis. **b** Heatmaps of TC-NER excision reads re-analyzed from XR-seq data of GG-NER deficient cells (XP-C) which are detected on non-coding + (blue) or – (red) strand of the genome, in genomic regions 2kb around TSSs for categories defined in **Fig. 3a-c** (see Methods). Dotted grey lines denotes the CAGE summits detected on each strand, and the dark blue dotted lines indicate positions 500 bases downstream of CAGE summits for the corresponding strand. **c** UCSC genome browser snapshots of representative loci for genomic categories defined in **b**. Tracks shown are CAGE (for + and – strand), Pol II-ser2P ChIP-seq (NO UV, +UV), TC-NER-specific XR-seq (for + and – strand). Coordinates of the genomic loci are shown at the top of each snapshot **d** Average profiles of read densities derived from **b**. For active bidirectional and unidirectional promoters only the divergent (DIV) loci were considered. **e** (Left panel) Scheme representing the window and strand taken in consideration for calculating Log_2_ FC between + strand and – strand at divergent loci. (Right panel) Box plots showing quantifications of the ratio of reads in between directions (indicated window sizes and borders) at bidirectional promoters for CAGE reads (shown in **Fig. 3**) and TC-NER-specific XR-seq reads shown in **b**. Box plots show the 25th– 75th percentiles, and error bars depict data range to the larger/ smaller value no further than 1.5 * IQR (inter-quartile range, or distance between the first and third quartiles). Two sample F-tests were conducted for each of 10,000 sampling pairs of 100 data points with replacement from each population, to test for significant difference between sample variance. The calculated P expresses the percentage of the non-significant F-tests (F-test p-value >= 0.05) out of the 10,000 total tests. **f** Comparison of CAGE and TC-NER-specific XR-seq reads as in **e** but between divergent non-overlapping mRNA and PROMPTs (indicated window sizes and borders). 95% confidence intervals (CI) of mean differences between log_2_ counts of tested conditions were calculated for 10,000 samplings of 100 data points with replacement from each population. Effect sizes of log_2_ counts between datasets were calculated using Cohen’s method (CES)

Next, we further investigated repair of PROMPTs and enhancers, a phenomenon previously observed, but hardly explained^24,53^. We quantified strand-specific repair upstream and downstream of unidirectional promoters and found stronger than expected repair activity at unambiguously resolved divergent PROMPTs (Fig. 6b-d, DIV). Indeed, XR-seq read density was not correlated to the steady state levels of CAGE at those loci (Pearson Correlation Coefficient = 0.1343). Also, FC of TC-NER reads between mRNA and PROMPTs were much smaller than for CAGE (Fig. 6f, 95 % CI excludes 0), thus matching the UV-independent Pol II-hypo uniformity (Supplementary Fig. 6 a-b). Similarly, TC-NER levels on TS of eRNAs were higher than anticipated. Indeed, density of eRNA XR-seq reads were similar to those of mRNAs (Fig. 6b-d) and contrasted with the very low CAGE signal detected at these loci (Fig. 3a-c). Therefore, balanced Pol II-hypo loading in PICs at all classes of transcripts allows for equal initiation events and mirrors the homogenous levels of XR-seq detected in these regions. Taken together, our results demonstrate that the widespread continual initiation and release into productive elongation of Pol II waves maximizes repair activity regardless of prior-to-UV transcript expression level at all kinds of active regulatory regions (mRNA, PROMPTs, enhancers).

### Continuous transcription initiation maximises transcription-driven repair

We next assessed the biological purpose of continuous transcription initiation from active regulatory regions during the UV-recovery period. We have reported previously^28^ that in the absence of a UV-triggered PPP release of elongating Pol II waves, the pri-elongating (e.g. already elongating prior to UV) Pol II molecules cannot repair the totality of the transcribed genome. Thus, we and others believe that sending Pol II molecules to allow the detection of the next lesions in line on the TS is of pivotal importance^28,42^.

To delineate this concept further, we quantified excision activity at thymidine dimers (TTs) (see Methods and^28^) as time passes using XR-seq data from XP-C cells irradiated under conditions that allow PPP-release of Pol II for only a short period of time (DRB2 experiment by^42^) (compare “DRB2 +0.5 h” with “DRB2 +1 h” in Supplementary Fig. 7 a-c). Critically we find, that the number of excision events in cluster I and II (more downstream of TSS) 1 hour after UV (DRB2 +1 h) did not match the levels of cluster 0 and I (more upstream) for DRB2 +0.5 h (before the asterisk in Supplementary Fig. 7c). Taking into consideration that only one Pol II molecule can be accommodated per PPP site at each active mRNA, PROMPT and eRNA allele at the time of irradiation, one can postulate that the extent of ongoing repair activity observed downstream of TSSs without DRB, 1 hour after UV, is the result of concurrent Pol II recruitment and initiation.

When we mapped XR-seq reads at TTs 1 h, 4 h and 8 h after UV in WT cells (data obtained from Adar et al.^53^, see Methods), we confirmed that significant levels of transcription-dependent excision activity was maintained for a significant proportion of lesions located directly downstream of active genes TSSs (cluster 0), even at late time points during the recovery process (Fig. 7a-c). Notably, it also appeared that the lesions located more distally to TSSs in active genes’ TS (from 32,5 kb to 1 Mb) get recognized and excised more efficiently as time passes (Fig. 7a-c, clusters III-IV and V-VI). These results reveal that a large extent of the transcription-driven repair activity is due to the ongoing entry of recycling Pol II molecules at TSSs. Our analysis highlights the advantage of a continuously supplied transcription-dependent repair process over slower and less efficient lesion detection capabilities of GG-NER, which was detected at significantly lower levels in all clusters as shown in inactive genes (Fig. 7 a-c).

**Fig. 7.**
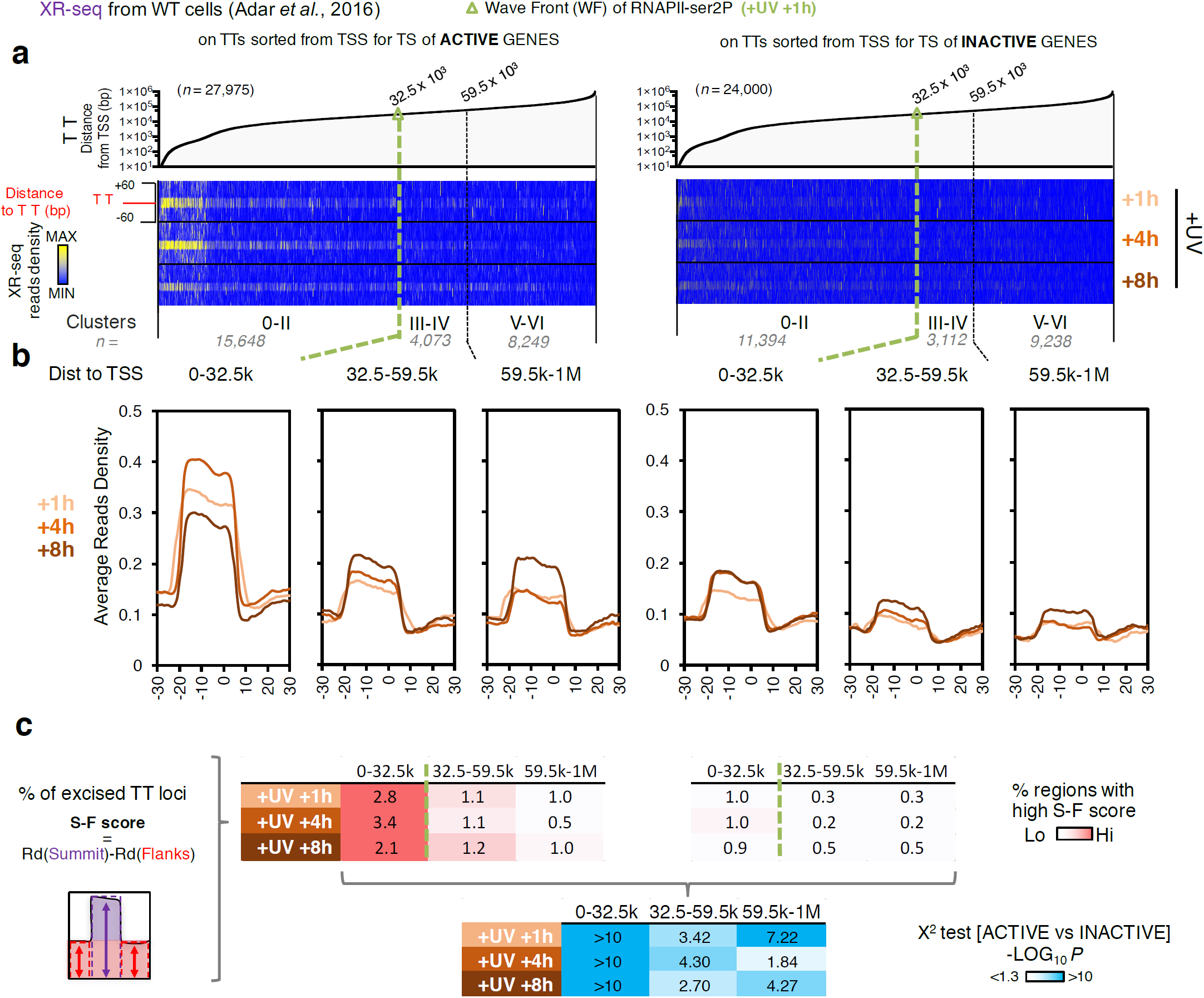
Continuous initiation maintains TC-NER efficiency over time and across the whole transcribed region **a** Distribution of XR-seq reads derived from WT cells at 1h, 4h and 8h post UV irradiation. Reads were aligned on TT loci of transcribed strand (TS) of active (left panel) and inactive (right panel) genes. Data were obtained from ^53^. Pol II-ser2P wave front position (WF =32.5 kb) was defined at +UV (+1h) in^28^. **B** Average plots of read densities showed in **a** for the respective clusters of active (right) and inactive (left) genes. **c** Plot showing the percentage (%) of excised TT loci, as calculated by the difference between Rd at summit (S) and Rd at flanks(F) (S-F score) from all clusters presented and analysed in **a** and **b**

## Discussion

In this study, we provide quantitative insights into the molecular processes underlying the major transcription-coordinated cellular response that is activated in human cells upon genotoxic stress^28–30,38,41,54^. The establishment of precise maps of chromatin state helped us to query in detail the impact of transcription on DNA repair activities at important functional regions, including PROMPT and eRNA loci. Our results support a model of continuous transcription initiation that can promptly feed the widespread UV-triggered escape of Pol II into the elongation enabling efficient DNA lesion-scanning of the whole transcribed genome.

The finding that increase in chromatin accessibility parallels the conservation of H3K27ac-modified nucleosomes at the flanks of already open regions in response to mild doses of UV-C irradiation is compatible with reports showing that there can be a significant gain in nucleosome accessibility without changes in nucleosome occupancy during rapid transcriptional induction^55^. Notably, the maintenance of H3K27ac at these sites prevents the imposition of repressive tri-methylation at active loci (see Fig. 2), in accord with the rule that H3K27ac and H3K27me3 are mutually exclusive^56^. Moreover, finding active transcription at these loci complies with prior reports suggesting that increase in gene expression are associated with surges in chromatin accessibility^57,58^ and that the presence of nRNA is known to inhibit^51,59^ the recruitment of H3K27me3-catalysing Polycomb Repressive complex 2 (PRC2) at active genes. Our data contrast the drastic chromatin remodeling observed in mice at a later time during recovery (6 h) when much higher doses of UV-B were used^60^. This suggests that when cells deal with unmanageable levels of damages they need to implement completely different expression changes required for the associated fate of programmed death, a protective mechanism limiting the risk of malignant transformation ^61–63^.

Our analysis takes advantage of a high-resolution strand-specific map of TSSs for coding and non-coding (enhancers and PROMPTs) loci and supports the idea that bidirectional transcription of divergent RNAs arises from two distinct hubs of transcription initiation (PICs), located within a single nucleosome-depleted region (NDR)^9,64–66^. Indeed, for bidirectional mRNAs and mRNA-PROMPTs, the binding of Pol II-hypo occurs at both edges of highly accessible regions (see ATAC-seq vs Pol II-hypo in Supplementary Fig. 6a, c), which correspond to single NDRs flanked by H3K27ac nucleosomes (see arrows in Supplementary Fig. 6c). These observations also extend the evidence supporting the claim that enhancers and PROMPTs PICs are organized in a similar manner to genes PICs^9,15^. In addition, the observed differences in transcript levels between PROMPTs and mRNAs (see Fig. 5) are probably not due to differences in Pol II-hypo recruitment (see Supplementary Fig. 6b, bottom), but rather due to differences in the frequency of premature termination at PPP sites and/or differences in degradation of PROMPTs RNAs by the RNA exosome. Interestingly, the latter is known to be inhibited upon UV stress ^67,68^ (see below).

By uncoupling TSSs of mRNA genes from those of PROMPTs and enhancers, we reveal that P-TEFb-dependent release of elongating Pol II from PPP sites extends to all actively transcribed regions (see Fig. 3). Interestingly, a growing number of studies have reported data to suggest that (i) UV irradiation preferentially inhibits elongation, rather than transcription initiation^28–30,38^, (ii) P-TEFb and NELF are important regulators of UV-response^41,54,69^ and (iii) although elongation gradually decelerates due to the encounter of Pol II with DNA lesions, significant initiation/early elongation activity (assessed by nRNA-seq) is observed in the first thousand bases of actively transcribed regions^28,29,38^, a characteristic that has also been used for the identification of active TSSs genome-wide after UV^30^. These features are consistent with our finding that new Pol II-hypo molecules are constantly recruited to PICs post-UV (see Fig. 4), and that they promptly proceed into initiation of start-RNAs and subsequently into elongation of longer nRNAs (see Fig. 5). Considering that Pol II ChIP density depends, among others, on the epitope residence-time at a given genomic locus^70^ and that Pol II molecules recruited in the PIC are readily phosphorylated^28,32,33^, we propose that the rapid exchange of Pol II isoforms after UV irradiation represents a perfectly plausible cause for the decreased ability to detect Pol II-hypo molecules at TSSs upon UV (see Fig.2, 4 and^28^). Such a model explains previously published data concerning the presence of (i) PIC/basal transcription factors in nuclear extracts^33^ or upstream of genes’ TSSs (TFIIB)^37^and (ii) nRNAs at the beginning of genes^28–30,38^ upon UV.

We note that excision fragments (from XR-seq) are distributed more homogeneously at sense (mRNA) and antisense (PROMPT) strands of unidirectional TSSs and at enhancers (Fig. 6) than it could be predicted from the CAGE levels. This finding reinforces the possibility that efficient repair at stable and unstable transcripts is primed by the uniform recruitment of Pol II-hypo at all classes of PICs in the steady state (see Supplementary Fig. 6). Remarkably, UV induces its continuous and uniform transition into initiation (see Fig. 4) and constantly feeds PPP release of DNA lesion-sensing Pol II into the transcriptome (see fig. 3 and 5). This concept was further validated by re-analysing an experiment mapping repair after inhibition of PPP-release, and thus *a fortiori* Pol II initiation^42^. We find a drastic impairment of CPD excision at TTs located on TS upstream of the previously calculated Pol II wave front (WF) (low XR-seq signal for DRB vs DMSO in supplementary Fig. 7d), and decrease in the percentage of excision of damaged TTs located in cluster 0-II (see Supplementary Fig. 7 a-c, DRB vs DMSO), thus suggesting that post-UV initiation is crucial for the repair of sequences located directly downstream of TSSs. Critically, uniform and continuous recruitment of Pol II at TSSs is further shown to maintain this process trough recovery, as ongoing TC-NER activity is persistently detected directly downstream of TSS at 4 and 8 hours after UV (see Fig. 7, TTs on active genes).

In the view of these results and the fact that continuous nRNA-seq signal has been detected uninterruptedly between 2 and 12 h post-UV^29^, we propose that maintaining initiation is necessary to allow for the sensing of the next lesion in line, thus maximizing the probability to repair quickly all lesions located on actively transcribed TS. Indeed, as time passes more lesions get to be removed further downstream (towards the end of long genes) (Fig. 7, cluster III to VI (32.5 kb to 1 Mb).

Increase in TC-NER at regulatory regions has also been observed in *E. Coli*^71^and is compatible with the idea that the act of antisense transcription exerts a meaningful biological function^72^ conserved through evolution. Indeed, these DNA sequences may serve as binding sites for transcription factors or encode target sites for RNA binding proteins, enabling accurate regulation of topologically associated mRNA genes^66,73^. Given the effect of DNA repair on the landscape of somatic mutations in cancer tissues^19,20^, surveillance of these vital sequences impacts on cell’s fitness. We propose that our model could account for the low levels of substitutions recently observed upstream of genes’ TSSs and around DNAse hyper-sensitive (DHS) sites^74–76^.

Recent advances in the field of transcription regulation point to the fact that activation of paused genes is mediated through switching from a premature termination state of Pol II at PPP sites to a processive elongation state^13,16,17^, implying that continuous initiation is required for fast transcriptional induction^16^. Our results, showing that persistent initiation guarantees a prolonged transcription-coupled NER, are functionally linked to the fact that DNA damage-triggered widespread PPP release of a given Pol II is sufficient to drive immediate initiation of the next Pol II (see Fig. 4 and 5, Supplementary Fig. 4 and 5). In other words, the clearance rate of Pol II from TSSs highly depends on PPP status. These findings are in favor to the emerging concept that Pol II pausing has an inhibitory effect on initiation^77–79^ and highlight how this mechanism can function across the whole transcriptome. At the same time, our results provide a novel physiological relevance to why cells could gain from firing initiation continuously, as balance between promoter-proximal termination and escape into elongation allows dynamic responses to stimuli.

## Supplementary Information

**Supplementary Figure 1.**
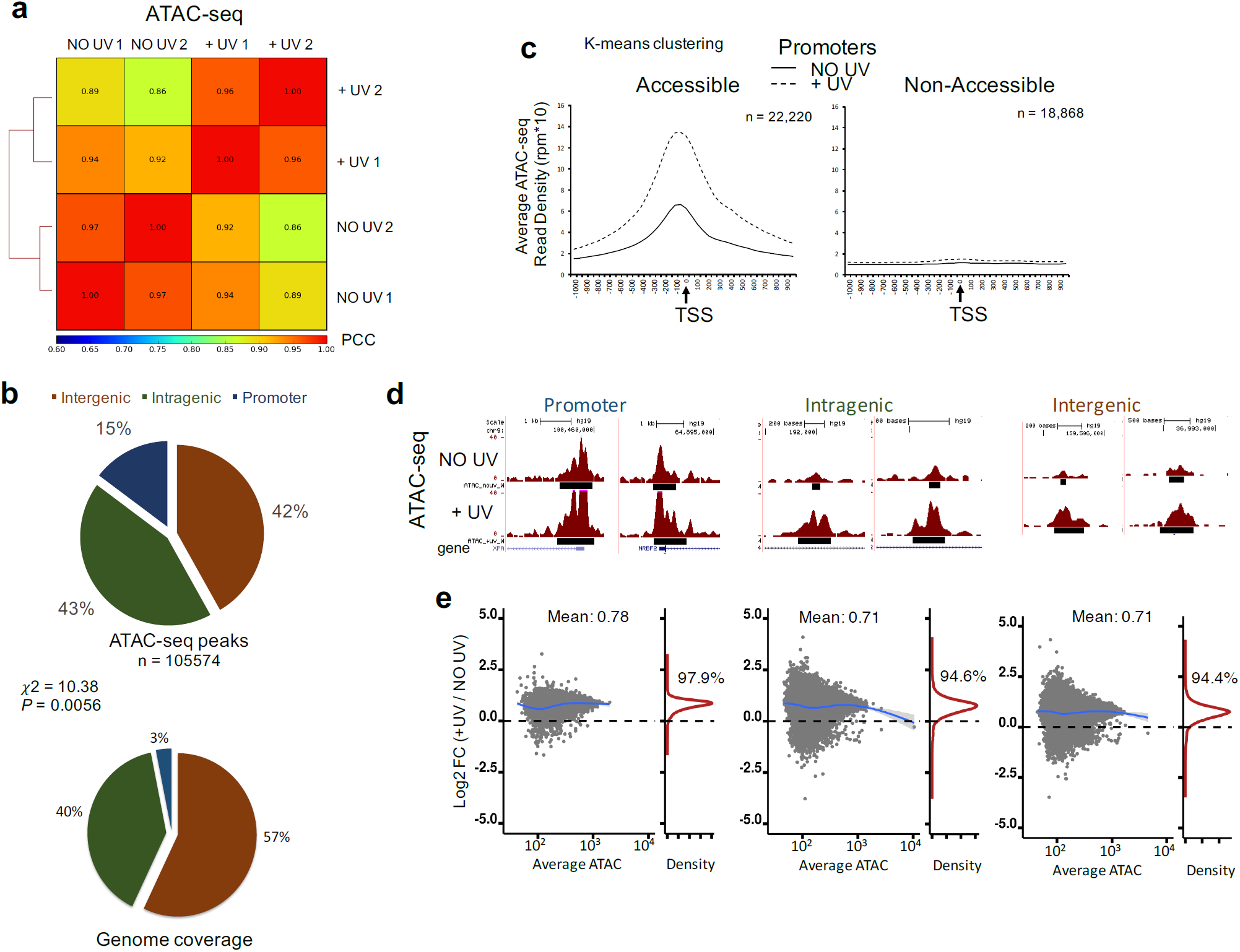
Increase of chromatin accessibility in response to mild doses of UV irradiation. **a** Correlation plot of ATAC-seq read densities for biological duplicates of non-irradiated (NO UV 1,2) and irradiated (+UV 1, 2) cells. Pearson’s Correlation Coefficients (PCC) were calculated genome wide (3kb windows) and reported on the heatmap. **b** Distribution of ATAC-seq peaks in promoter (+/-2kb relative to TSS), intragenic and intergenic regions (upper part), compared to the genome percentage covered by the respective regions (lower part). Chi-square test were performed to determine whether observed values differ from expected value purely by chance. **c** Average plots depicting ATAC-seq read densities in k-means resolved clusters of accessible and non-accessible promoters for non-irradiated (NO UV, solid line) and irradiated (+UV, dashed line) cells. **d** UCSC snapshot of ATAC-seq profiles of representative loci showing signal profile before (upper part) and after (lower part) UV, in promoter, intragenic and intergenic regions. **e** MA plots showing the individual (grey dots) and average (blue line) FC (Log_2_ FC) in ATAC read density at ATAC-seq peaks, between +UV and NO UV, as a function of the average (from replicates) ATAC-seq read density in NO UV. Percentage of peaks with increased FC (Log_2_ FC > 0) is indicated on the kernel density plot on the right of each MA plot.

**Supplementary Figure 2.**
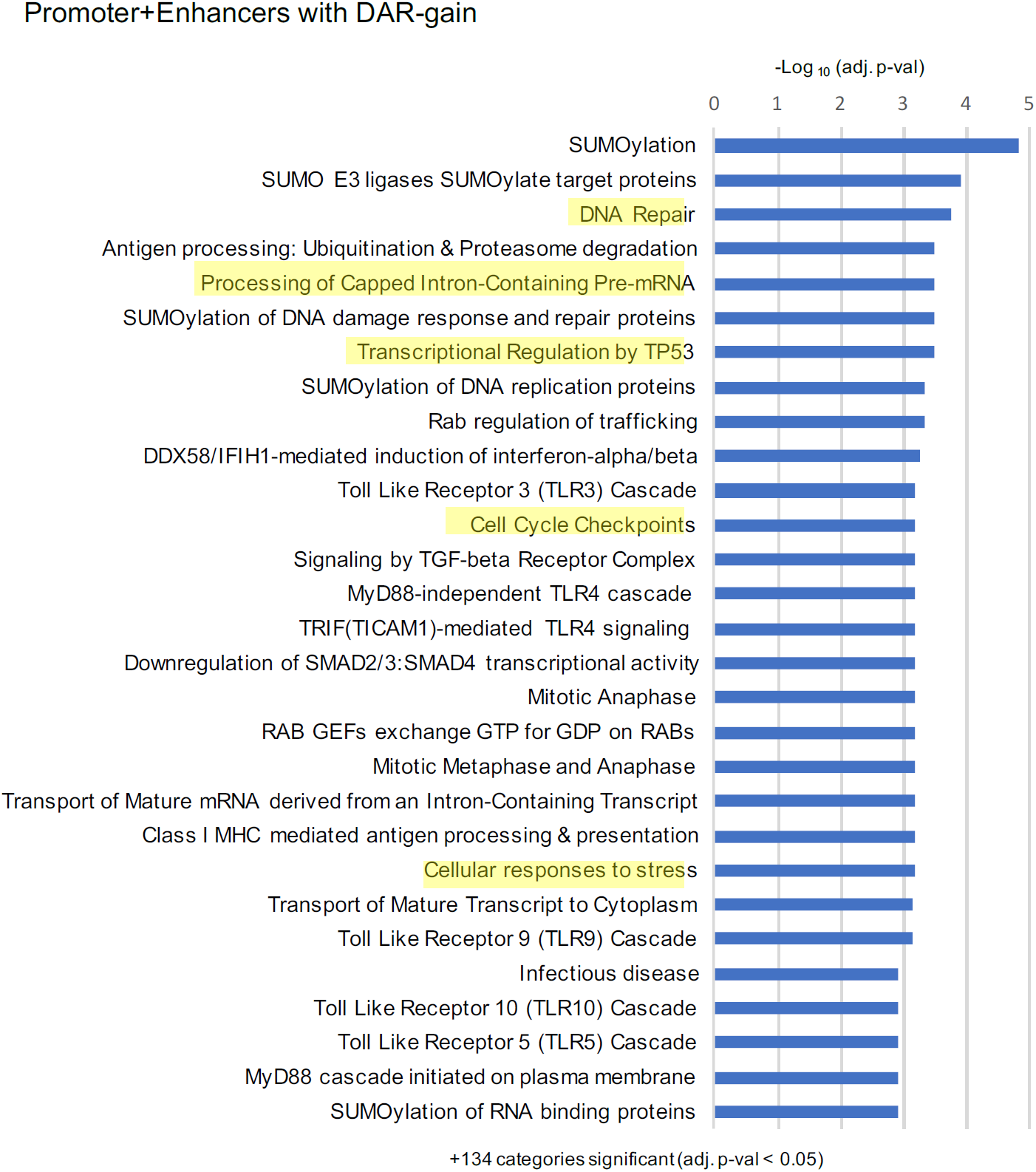
Gene Ontology analysis of DAR-gain regions. Reactome biological pathway (see Methods) analysis for genes that are associated with DAR-gain genomic regions (either proximally associated with promoter DAR-gain or controlled by a intergenic/intragenic DAR-gain region annotated as enhancer and associated topologically to a given gene according to FANTOM5 consortium). Pathways are sorted by decreasing significance. Biological pathways highlighted in yellow are relevant to the processes involved in UV-response or transcription

**Supplementary Figure 3.**
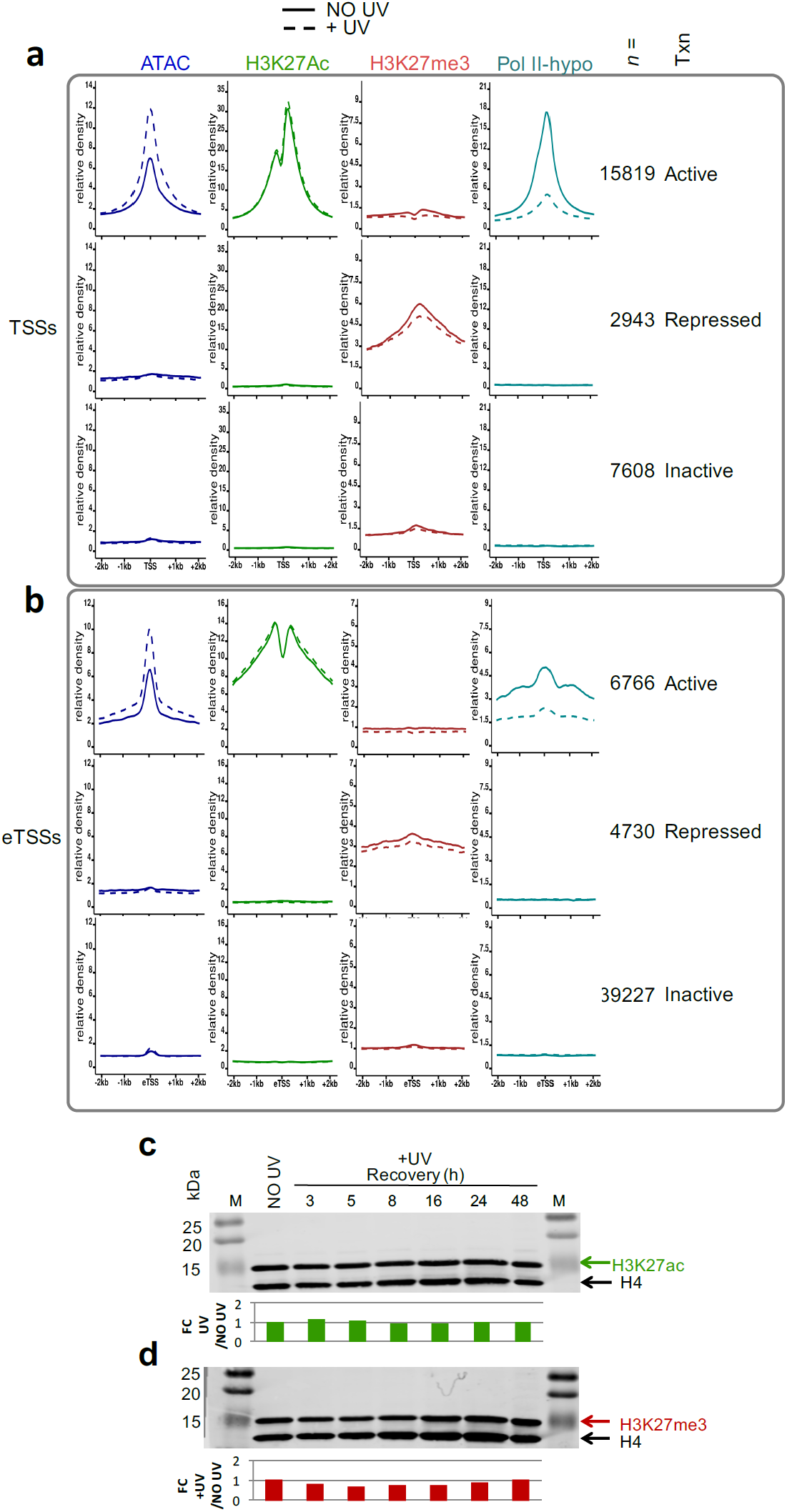
Chromatin modifications remain virtually stable upon UV damage. **a**,**b** Average profile plots illustrating the read densities for ATAC-seq, H3K27ac, H3K27me3 and Pol II-hypo ChIP-seq before (NO UV, solid line) and after UV (+UV, dashed line), on active, inactive and repressed TSSs **(a)** and eTSSs **(b)**. Data for Pol II-hypo are obtained from^28^. **c** Western Blot analysis of histone extracts for bulk H3K27ac levels before and after UV. Recovery times after UV are as indicated. Total histone 4 (H4) was used as a loading control. Quantification was calculated and compared to NO UV condition. The figure is representative of 2 independent experiments. **d** Same as in **(c)** but for H3K27me3

**Supplementary Figure 4.**
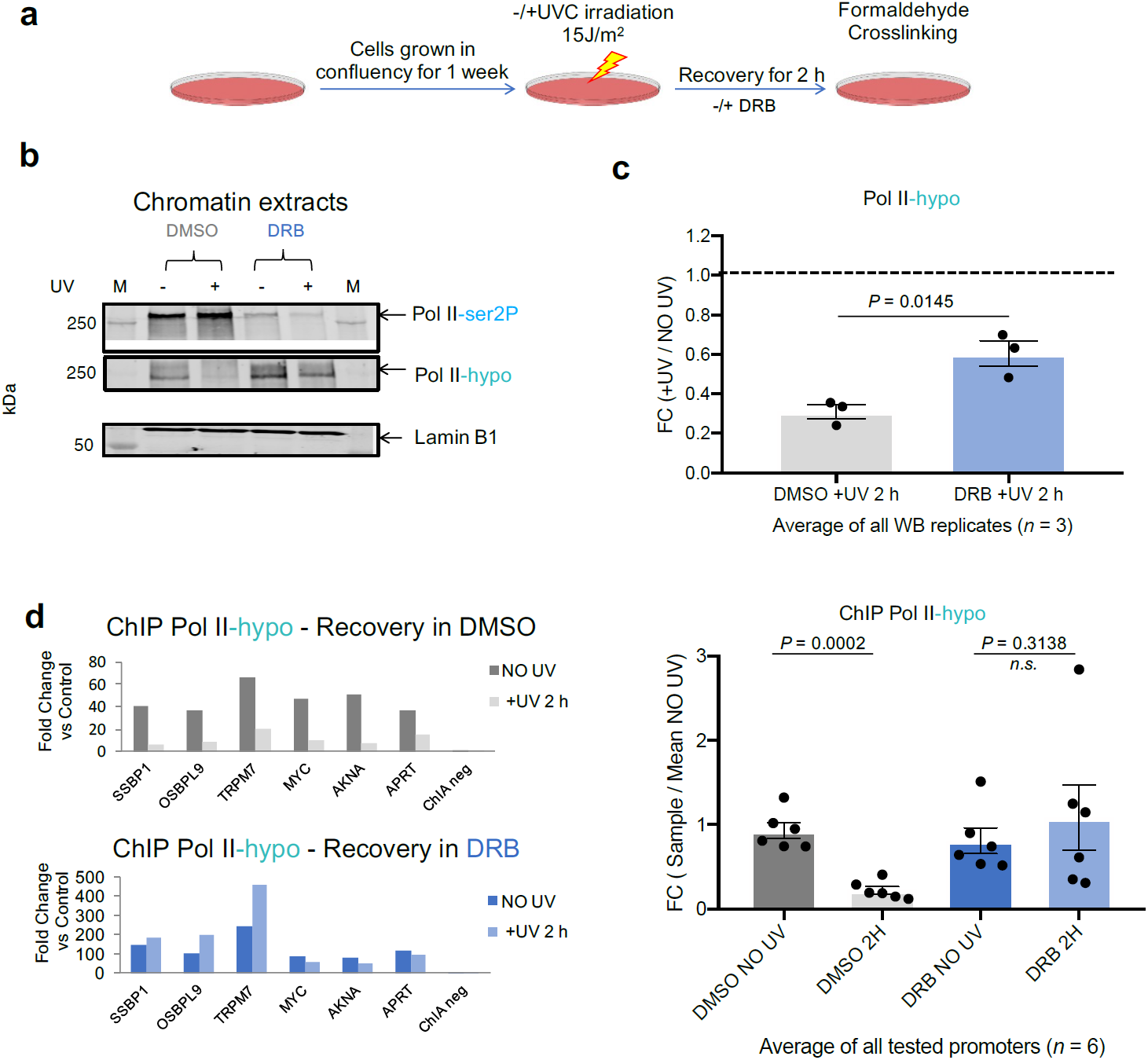
Rescuing levels of pre-initiating Pol II (Pol II-hypo) after UV irradiation. **a** Experimental timeline. DRB was added in a final concentration of 100 μM. **b** Western Blot analysis of chromatin extracts for Pol II-ser2P and Pol II-hypo in the experimental conditions outlined in **(a)**. Lamin B1 was used as loading control. The figure is representative of 3 independent experiments. **c** Quantification of the average signal detected in **(b)** as compared to NO UV (dashed line) condition for DMSO and DRB treated cells, respectively. Error bars represent S.E.M. and *P* values are calculated using two-sided Student’s t test. **d** (Left) ChIP-qPCR analysis of Pol II-hypo enrichment at the promoter regions of 6 representative genes selected from **Fig. 3** for cells treated as indicated in (a) with DMSO or with DRB. FC was calculated against a negative locus (ChIA neg). (Right) Average ChIP-qPCR enrichment of all genes tested (error bars represent +/-SEM). Error bars represent S.E.M. and *P* values are calculated using two-sided Student’s t test

**Supplementary Figure 5.**
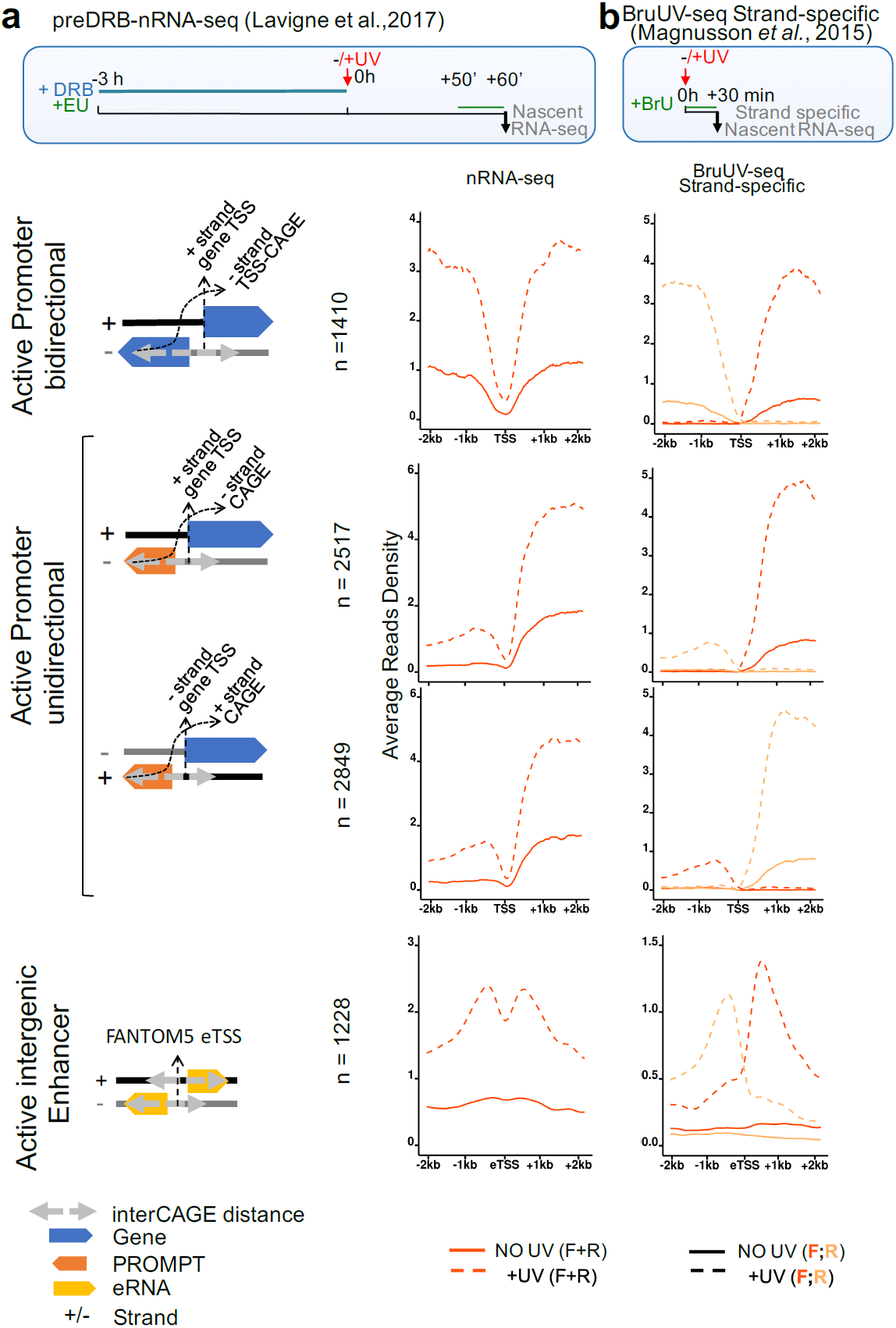
*De novo* and increased RNA synthesis from all TSSs upon UV exposure. **a** (Upper part) Scheme for preDRB-nRNA-seq methodology (see^28^). (Lower part) Average Read densities profiles of nRNA-seq (data from^28^, see Methods), before (solid line) and after UV (dashed line) in genomic regions 2kb for categories defined in Fig. 3a. **b** Same as in **(a)** for strand-specific protocol of BruUV-seq (see^30^)

**Supplementary Figure 6.**
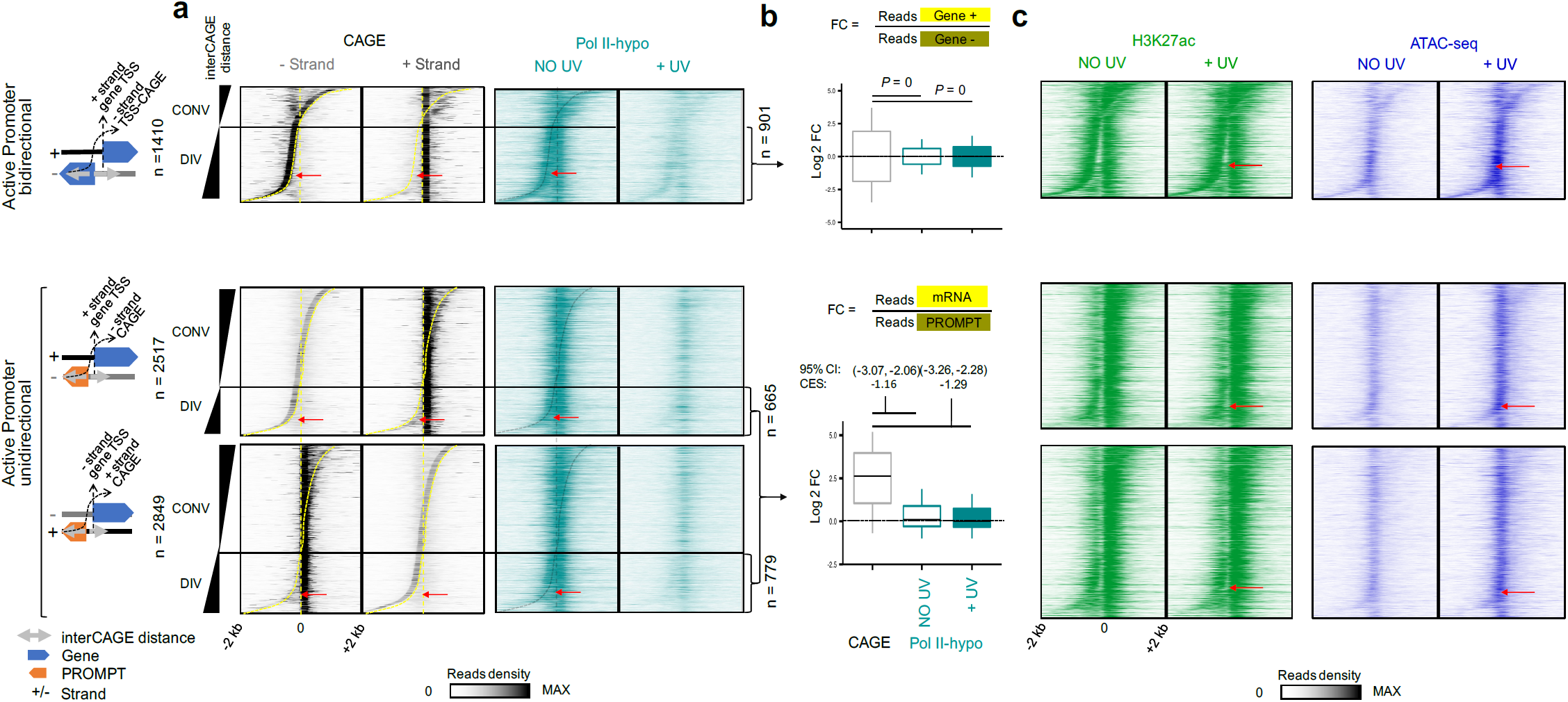
Equal recruitment of Pol II on the flanks of NDR primes for increased transcription initiation from gene TSS and divergent PROMPTs upon UV. **a** Heatmaps of read densities of CAGE (- and + strand) and Pol II-hypo ChIP-seq (data obtained from^28^, see Methods) at steady state condition (NO UV) and upon UV, in genomic regions extending 2 kb around TSSs for categories defined in Fig. 3a**. b** Box plots showing quantifications for CAGE and Pol II-hypo ChIP-seq reads showed in **(a)** of the ratio between (+ strand) over (-strand) for divergent bidirectional promoters (upper) and mRNA over PROMPT (lower) for divergent unidirectional promoters. Box plots show the 25th– 75th percentiles, and error bars depict data range to the larger/ smaller value no further than 1.5 * IQR (inter-quartile range, or distance between the first and third quartiles). (Upper) Two sample F-tests were conducted for each of 10,000 sampling pairs of 100 data points with replacement from each population, to test for significant difference between sample variance. The calculated P expresses the percentage of the non-significant F-tests (F-test p-value >= 0.05) out of the 10,000 total tests. (Bottom) 95 % confidence intervals (CI) of mean differences between log_2_ counts of tested conditions were calculated for 10,000 samplings of 100 data points with replacement from each population. Effect sizes of log_2_ counts between datasets were calculated using Cohen’s method (CES). **c** Same as in **(a)** for Pol II-hypo but for H3K27ac ChIP-seq (green) and ATAC-seq (blue) reads. Red arrows indicate the Nucleosome Depleted Regions (NDRs) for active divergent bidirectional and unidirectional promoters

**Supplementary Figure 7.**
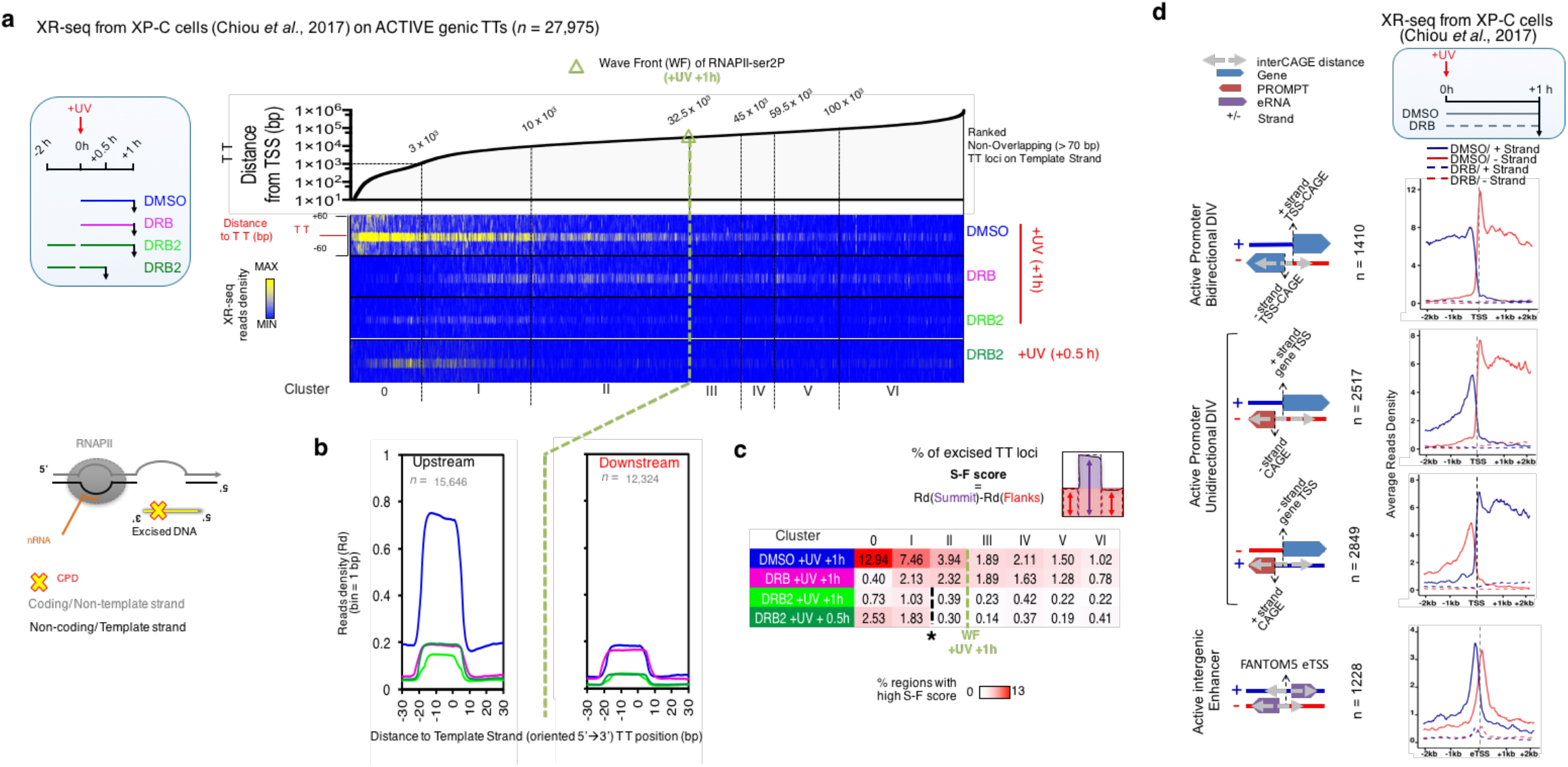
TC-NER activity in sets of DRB treated conditions reveals the need for continuous transcription initiation for repair of the totality of the active transcriptome. **a** Heatmap depicting the distribution of excised DNA fragments (XR-seq) derived from TC-NER activity (XP-C cells). Reads were aligned around TT loci of transcribed strand, localized on active genes. Data were obtained from^42^. Box in blue border (left) illustrates the experimental timeline followed and the respective drug treatments. **b** Average plots of read densities (Rd) showed in **a** but for clusters upstream (0, I, II) and downstream (III, IV, V, VI) of Pol II-ser2P wave front (WF) as defined in^28^ for +UV (+1h) condition. **c** Plot showing the percentage (%) of excised TT loci, as calculated by the difference between Rd at summit (S) and Rd at flanks(F) (S-F score) from all clusters presented and analyzed in **a** and **b. d** Average profile of read densities of XR-seq signal derived from XP-C cells. Dashed lines illustrate the XR-seq signal in + (blue) or – (red) strand, in the presence of DRB.

## Materials and Methods

### Cell culture and treatments

Cells used in this study were VH10 HTERT immortalized normal human skin fibroblasts. Cells were cultured, synchronized by low-serum-starvation and release in full medium as described previously^28^, unless stated differently. When applied, 5,6-dichloro-1-β-D-ribofuranosylbenzimidazole (called DRB) (Calbiochem) and triptolide (called TRP, Invivogen) were used in a final concentration of 100 μM and 125 nM, respectively and they were added directly in growth media at indicated times. Cells were irradiated with mild doses of UV-C (254 nm, TUV Lamp, Philips) (15 J/m^2^ except if otherwise stated).

### ChIP-seq

ChIP-seq was performed as previously described^28^ with minor changes. Cells were mock-treated (NO UV) or with UV (+UV) (Fig. 2, dose for H3K27me3 was 20 J/m^2^) or as indicated on the timeline (Fig. 4). Formaldehyde was added to the cells at a final concentration of 1% at 4°C for 12 mins. The reaction was quenched by adding glycine (final concentration of 125 mM) for 5 mins. Cells were washed 3 times with cold PBS and then collected in PBS containing 1 mM EDTA, 0.5 mM EGTA and 1 mM PMSF and pelleted in pellets of ∼2×10^7^ cross-linked cells.

Pellets were resuspended in Chro-IP lysis Buffer (50 mM Hepes-KOH pH 8.0, 1 mM EDTA, 0.5 mM EGTA, 140 mM NaCl, 10 % glycerol, 0.5 % IGEPAL, 0.25 % Triton X-100, 1 mM PMSF, and a mix of protease inhibitors (Roche)). After 10 min rotation at 4°C, cell suspension was centrifuged (10 min, 2,800 rpm at 4°C) and supernatant was kept as soluble fraction. In turn, cell pellet was washed with Wash Buffer (10 mM Tris-HCl, pH 8.0, 1 mM EDTA, 0.5 mM EGTA, 200 mM NaCl, 1 mM PMSF, 10 mM NaPy and protease inhibitors). The cell suspension was rotated for 10 min at 4°C and centrifuged as mentioned above. The cell pellet was resuspended in RIPA Buffer (10 mM Tris-HCl, pH 8.0, 1 mM EDTA, 0.5 mM EGTA, 140 mM NaCl, 1% Triton X-100, 0.1% Na-Deoxycholate, 0.1% SDS, 1 mM PMSF, 10 mM NaPy and protease inhibitors). Samples were sonicated using the Bioruptor water bath sonicator (Diagenode) using the “high” setting with cycles of 30 sec “on” and 30 sec “off”, for a total duration of 25 minutes. Samples were then centrifuged for 10 min at 10,000 rpm at 4°C. The supernatant was kept as chromatin fraction (Input). Chromatin immunoprecipitation (ChIP) was performed by incubation of equal amounts of sheared chromatin from irradiated and non-irradiated cells with the appropriate antibody at 4°C overnight. The antibodies used for ChIP were the following: H3K27ac (ab4729, Abcam), H3K27me3 (07-449, Millipore), 8WG16 (Pol II-hypo) (05-952, Millipore). Protein A (for H3K27ac and H3K27me3) or Protein G (for 8WG16) Dynabeads (ThermoFisher Scientific) were blocked overnight at 4°C in RIPA buffer without protease inhibitors, NaPy and PMSF in the presence of Bovine Serum Albumin (BSA) (30μg/ml). Next day, beads and immunoprecipitated chromatin were co-incubated for 3 h at 4°C and then beads were sequentially washed twice with RIPA, three times with RIPA containing 0.3M NaCl, once with LiCl buffer (0,25M LiCl, 10mM Tris-HCl PH 8.0, 1mM EDTA, 0.5mM EGTA, 0.5% Triton X-100, 0.5% Sodium deoxycholate) and twice with TE buffer (10 mM Tris-HCl PH 8.0, 1 mM EDTA pH 8.0). Chromatin immuno-complexes were eluted by two rounds of incubation at 65°C for 20min in 1% SDS and 100 mM NaHCO3, and vigorous vortexing. Finally, de-crosslinking of Input and immunoprecipitated chromatin was performed by overnight incubation at 65°C in the presence of 200 mM of NACl. We then applied Proteinase K treatment (0.1μg/μl in 0.5% SDS) for 1h at 55°C, and DNA purification was performed with AMPURE XP Beads (Agencourt) according to manufacturer’s protocol.

The primers used for ChIP-qPCR experiments were the following (5’ to 3’, F:Forward, R:Reverse, ChIA neg was the negative primer): SSBP1_F: GTGAGGGAGGAAGGGATAGC, SSBP1_R: AGGGCCAGACACCTACACAG, OSBPL9_F:ATTGGCGGCTCCCAAGAT, OSBPL9_R: GCATTGTAGTCCAGCACGAA, *TRPM7_*F: CCCAGGGAAACCTTCTCAG, TRPM7_R: TCGCACAATTATGAAAGACTCG, *MYC*_F: ACTCAGTCTGGGTGGAAGGTATC, *MYC*_R:GGAGGAATGATAGAGGCATAAGGAG, AKNA_F: CCGTTCCAATCCCTTACC, AKNA_R: TGGAACAAAGAATTCACAGG, APRT_F: GCCTTGACTCGCACTTTTGT, APRT_R: TAGGCGCCATCGATTTTAAG, ChIA_neg_F: AGTCTGAGCTTTGTGGACAGC, ChIA_neg_R: CCCTCCCAGTATACAGTCTTGC. qPCR, library preparation and next-generation sequencing were performed as previously described^28^.

### Histone acetic extraction

Cells grown to confluence in 10 cm plates, synchronized and released as described above were irradiated with 15 J/m^2^. Cells were washed 3 times with cold PBS, harvested and centrifuged at 2000 rpm for 5 min at different recovery periods as indicated in Supplementary Fig. 3c, d. Supernatant was discarded and cell pellet was washed with 10 volumes of cold PBS and centrifuged at 2000 rpm for 5 min. Supernatant was removed and cell pellet was suspended in 10 volumes of Lysis Buffer (10 mM Hepes pH 7.9, 1.5 mM MgCl_2_, 10 mM KCl, 0.5 mM DTT, 1.5 mM PMSF), sulfuric acid was added in a final concentration of 0.2M and the suspension was incubated on ice for 30 min before being centrifuged at 10,080g for 10 min at 4°C. Next, the supernatant fraction was collected and TCA was added in a final concentration of 20%. Samples were vortexed and kept on ice for 1 h. Next, samples were centrifuged for 15 min at 14,000 rpm at 4°C. Supernatant was discarded and the pellet was washed with 1 ml of ice cold (−20°C) acetone. After centrifugation at 14,000 rpm at 4°C for 5 min, acetone was removed carefully using a centrifugal evaporator and pellet was resuspended in TE buffer and stored at −80°C.

### Western Blot analysis

Western Blot analysis of equal amounts of crosslinked chromatin extracts or equal amounts of histone extracts was performed as described^28^. Antibodies used for Western Blot analysis are the following: anti-H3K27ac (ab4729, Abcam), anti-H3K27me3 (07-449, Millipore), 8WG16 (05-952, Millipore), anti-elongating RNA pol II (ab5095, Abcam), anti-Lamin B1 (ab65986, Abcam), anti-histone 4 (ab10158, Abcam), anti-histone3 (ab1791, Abcam). Time for analysis are indicated on the Figures (Fig. 4b and Supplementary Fig. 3 c, d).

### Assay for Transposase Accessible Chromatin (ATAC)-seq

ATAC-seq method (nuclei preparation, transposition and amplification of transposed fragments for library preparation) was performed using Nextera DNA Library Prep Kit (Illumina, Inc) and primers as described in Corces et al.^43^ with minor modifications; (i) 70,000 cells were used per experimental condition and (ii) The DNase treatment of cells in culture medium, before the transposition reaction, was skipped. The UV dose applied for ATAC-seq experiments was 15 J/m^2^ and treated cells were left to recover for 2 h before harvesting.

### Start-RNAs isolation and qPCRs

To isolate small RNAs (smaller than 200 nucleotides), we used Qiagen miRNeasy Mini Kit and RNeasy MinElute Cleanup Kit according to manufacturer instructions. In order to monitor the efficiency of the different enzymatic reactions, we included in our experiments a spike-in RNA oligonucleotide of known sequence (oGAB11: rArGrUrCrArCrUrUrArGrCrGrArUrGrUrArCrArCrUrGrArCrUrGrUrG, synthesized and purified by IDT). After purification, small RNAs and spike-in molecules were ligated to the IDT DNA linker 1 (/5rApp/CTGTAGGCACCATCAAT/3ddC/). Specifically, samples were denatured for 2 mins at 80°C and then placed immediately on ice. Ligation mix (4,8 μl 50% PEG, 2 μl 10x RNA ligase Buffer, linker and RNase free H20, 0.5 μl truncated RNA ligase (NEB, Cat No. M0351S) was added in a final volume of 20 μl. The reaction was incubated for 3 h at 37°C. After H20 was added to a final volume of 100μl, ethanol precipitation (3 volumes of 100 % EtOH, with 1/10th volume of 3M NaAc, pH 5.2, and 10 μg of Glycogen (ThermoFischer Scientific, Cat Number AM9510) was performed overnight at −80°C. RNA was purified in 10 μl and Reverse Transcription (RT) was performed using primer oLSC003: /5Phos/TCGTATGCCGTCTTCTGCTTG/iSp18/CACTCA/iSp18/AATGATACGGCGACCACCGATCCGACGATCATTGATGGTGCCTACAG according to Invitrogen Superscript II (Cat Number 18064014) instructions. qPCR was performed using gene specific forward primers (Sequences 5’ to 3’ for *OSBPL9*: ATTGGCGGCTCCCAAGAT, *SSBP1*: GTGAGGGAGGAAGGGATAGC, *IFIT1*: TCTCAGAGGAGCCTGGCTAA, *KPNA6*: ATTTGGCGAGAGCCTGTCT) and one common reverse primer (oNTI230: 5’-AATGATACGGCGACCACCGA-3’), which anneals to RT primer oLSC003 sequence.

### Read alignment, normalization, peak calling and differential accessibility analysis

For all Next Generation Sequencing (NGS) data analyses, in-house scripts and pipelines were developed to automate and analyze the data consistently (see below for details). Code is available upon request. Sequenced data and generated wig profiles are available on Gene Expression Omnibus (GEO) (Accession ID: GSE125181). Short read quality control, data filtering alignment and wig profile generation was performed essentially as described previously^28^ with minor modifications.

Chip-seq data for pol II-ser2P and pol II-hypo (Fig. 2,3 and Supplementary Fig. S3,6), were obtained from^28^ for NO UV and 2 h or 1.5 h post-UV (8 J/m^2^), respectively. nRNA-seq data were obtained from^28,30^ and processed as described in^28^ (Fig. 5 and Supplementary Fig. 5). CAGE alignments were obtained from FANTOM5 (see below).

For H3K27ac and H3K27me3 ChIP-seq alignment files, peak calling was performed using SICER version 1.1^80^ with window parameter = 400 bp and gap parameter = 1, while fdr and *log*_*2*_fold change cutoffs were set to 0.01 and 1.5 respectively. For ATAC-seq alignment files, peak calling was performed using MACS2^81^. Because of the variability of ATAC-seq fragment lengths, several runs of the peak calling algorithm were performed, using different parameters per run, in an attempt to maximize the sensitivity of the detection of open chromatin regions. In particular, --nomodel --shift 100 --extsize 200, --broad --nomodel --shift 100 --extsize 200 --keep-dup all, --nomodel --shift 37 --extsize 73, --broad --nomodel --shift 37 --extsize 73 --keep-dup all, --nomodel --shift 75 --extsize 150 --keep-dup all runs were combined, and detected peaks were filtered using fdr < 0.05 and fold change > 1. To perform differential accessibility analysis, diffBind R package (https://www.bioconductor.org/packages//2.10/bioc/html/DiffBind.html) was used, with the combined peak sets as a consensus open chromatin reference. Differential accessibility regions were detected and filtered by applying fold change (Log_2_ FC ≥ 1) and p-value (p-val ≤ 0.001) thresholds.

### Read density plots

ATAC-seq, ChIP-seq, nRNA-seq, CAGE-seq and XR-seq data were subjected to read density analysis after read depth normalisation of all samples *per* experiment. Heatmaps and average density profiling were computed as previously^28^ around genomic regions of interest, as indicated in the figures. Heatmaps were generated directly using the software, from matrices of binned read densities (bin size is indicated in the figures) for all considered individual (n) items (metagenes). Read density matrices were also imported in R and python custom scripts for (i) plotting average density profiles (smoothing achieved by a moving window of the bin size as indicated) and (ii) for determination of read densities per genomic category.

### Construction of mRNA-TSSs, PROMPT-TSSs and eTSS annotation

To annotate Transcription Start Sites (TSSs), all known protein coding and non-coding RNA hg19 RefSeq transcripts release 86 were downloaded from UCSC table browser (http://genome-euro.ucsc.edu/cgi-bin/hgTables). For each transcript, a biotype was assigned using BioMart (www.biomart.org), and all the small non-coding RNAs were excluded. For all the gene models containing multiple alternative transcripts, TSS neighborhoods of a 100 bp window were clustered together, and only the longest transcript was kept, resulting to 30,473 transcripts. Transcripts were then separated into 3 groups, based on their transcriptional activity. TSS coordinates were extended to 2kb on each direction and were tested for overlap with the Pol II-ser2P -UV, H3K27ac -UV and H3K27me3 -UV peak sets. Regions overlapping with an Pol II-ser2P -UV and H3K27ac -UV peak, were characterized as active, those overlapping with an H3K27me3 -UV peak, but not with an Pol II-ser2P -UV neither with a H3K27ac -UV peak were characterized as repressed, and those that did not overlap with any of the above peak sets were characterized as inactive. Any region overlapping with both H3K27ac -UV and H3K27me3 -UV peaks were excluded from the rest of the analysis. This resulted to 15,819 active, 2,943 repressed and 7,608 inactive transcripts. To further classify the active TSSs in terms of transcription directionality, the annotation was split up into unidirectional and bidirectional references. All active transcript pairs with opposite direction of transcription, where −2 kb ≤ TSS_distance_ ≤ +2 kb, TSS_distance_ = TSS _coordinate_forward strand_ –TSS_coordinate reverse strand_ (interCAGE distance) were characterized as bidirectional, while the rest of the annotations were characterized as unidirectional. Bidirectional pairs were further categorized into two groups of annotations: convergent bidirectional transcript pairs with TSS_distance_ ≤ 100 bp, and divergent bidirectional transcript pairs with TSS_distance_ > 100 bp. To optimize the categorization of convergent and divergent transcript pairs, TSS coordinates were redefined by scanning in a radius of 250 bp, to detect the nucleotide occupied by the maximum sense CAGE signal. Any bidirectional pair with a non-significant CAGE peak in the aforementioned region was excluded from the analysis. This finally resulted to 12,859 unidirectional transcripts and 2,822 active bidirectional TSS pairs, 1,806 of which were characterized as divergent and 1,016 as convergent.

To gain a complete overview of the non-coding antisense transcription events occurring around mRNA TSSs, we also annotated upstream antisense (uaRNA) and downstream antisense (daRNA) transcripts (referred as an ensemble to PROMPTs in this paper for convenience). Only the active unidirectional mRNA TSSs were used. For all the genes annotated with more than one mRNA transcript, only the leftmost TSS (for + strand genes), and rightmost TSS (for – strand genes) were considered for the rest of the analysis. The antisense CAGE peak with the highest summit in the region ranged from −2 kb upstream to +1 kb downstream of each unidirectional TSS was considered to be the main PROMPT TSS for further analyses. The above procedure was also repeated for the inactive transcript set, to estimate the highest CAGE summit background distribution. The putative active PROMPT CAGE summits, which were higher than the average of the summit background distribution, were considered as active. This resulted to 5,366 pairs of active unidirectional – PROMPT TSSs, which were categorized to 1,444 divergent and 3,922 convergent pairs, as described above. By focusing on the divergent loci, the dynamics of transcription could be studied at play in each direction, without having to deal with interference from either direction. Therefore, analysis was focused on upstream antisense RNA, which correspond to the original definition of PROMPTs^6^.

To annotate enhancer Transcription Start Sites (eTSSs), all 65,423 human enhancers from phases 1 and 2 of the FANTOM5 project from http://fantom.gsc.riken.jp/5/datafiles/phase2.2/extra/Enhancers/human_permissive_enhancers_phase_1_and_2.bed.gz, and the center of each annotation was considered as the corresponding transcription start site. Enhancers were separated to 6,766 active, 4,730 repressed and 39,227 inactive following the pipeline described above. Active intergenic enhancers were further analyzed, and all the eTSSs within a distance of 10kb from nearby active transcripts, or neighbor eTSSs within a distance of 2kb were excluded. The rest of the intergenic eTSSs were extended to 1kb in both directions, and sense and antisense maximum CAGE summit heights were detected for each reference. This procedure was also repeated for the inactive enhancer set and inactive sense and antisense highest CAGE summit background distributions were estimated as described above. Finally, the putative active intergenic sense and antisense CAGE summits which were higher the averages of the summit background distributions, were considered as active. This resulted to 1,228 active intergenic eTSSs.

### Promoter Escape Indices analysis

Promoter escape analysis was performed for a subset of active unidirectional and bidirectional transcripts, PROMPTs and active enhancers. In particular, to avoid the inclusion of Pol II-ser2P reads mapped in overlapping promoters and gene bodies, only active divergent unidirectional transcript – PROMPT pairs were considered, where TSS_distance_ > 100 bp, TSS_distance_= TSS _coordinate forward reference_-TSS _coordinate reverse reference_, active divergent bidirectional transcript pairs with TSS_distance_ > 100 bp and active intergenic enhancers with no nearby transcripts within 10 kb and no nearby eTSSs within 2kb. For TSSs and PROMPT-TSSs promoter Escape Indexes (EI) were calculated as previously defined^28^, by taking the average coverage in rpm in the gene body (density in gene body was abbreviated as Db and ranged from 101 bp to 2 kb downstream of TSS or 101 bp downstream of TSS to TTS for genes larger or smaller than 2 kb respectively) divided by the average coverage on the promoter-proximal region (Dp) ranged from 250 bp upstream to 100 bp downstream of TSS.

For enhancer escape analysis, EI was calculated as above, where Density of reads at enhancer flanks (Df) is calculated for the regions ranging from −2 kb to −100 bp upstream of eTSS and from +100 bp to +2 kb downstream of eTSS, while density of reads on enhancer TSS (De) is calculated for the regions ranging from 100 bp upstream to 100 bp downstream of eTSS.

### Estimation of the proportion of the inhibited transcriptome upon UV irradiation

For calculating the percentage of the normally transcribed genome showing transcription inhibition previously published nRNA-seq data (NO UV and +UV 2h from^28^) were used to determine the actively transcribed regions where signal ratio (Log_2_ FC (+UV/NOUV)) < 0. All active transcripts of length over 100kb were trimmed up to 100kb and were divided to genomic bins of 1kb. Read-depth normalized and exon-free nRNA-seq reads of each of the two conditions were counted on each genomic bin of each transcript, and for each of the n{1,2,…,100} bin positions the average Log_2_ FC (+UV/NOUV)) ratio was calculated for the set of the active transcripts. This resulted to a vector of size 100, with Log_2_ FC (+UV/NOUV)) values >= 1 for the first 28 bins, implying transcription clearance on the first 28kb of the active transcriptome upon UV damage, while for the last 72 bins Log_2_ FC (+UV/NOUV)) values < 0, implying transcription inhibition upon UV damage. To calculate the total proportion of the active transcriptome where transcription was inhibited, the coverage (in bp) of all the normally actively transcribed elements (see above for definition of these loci) located within 28 kb from TSS were summed up and divided by the total length of all the actively transcribed elements, resulting to 63.65 %.

### Nucleotide Excision Repair data meta-analysis

The strand-specific genome-wide maps of nucleotide excision repair of the UV-induced DNA damage (CPDs), available for XP-C mutants lacking the global-genome nucleotide excision repair mechanism (GG-NER-deficient, TC-NER-proficient) were obtained from Hu et al. ^24^ (data for Fig.6, Gene Expression Omnibus, (GEO) accession number GSE67941) and Chiou et al.^42^, (data for Supplementary Fig. 7, GEO accession number GSE106823). XR-seq data for wild type cells (used for Fig. 7, GEO accession number GSE76391) were obtained from Adar et al.^53^. Sequence Read Archive (SRA) datasets were downloaded from Gene Expression Omnibus using the sra toolkit prefetch (https://www.ncbi.nlm.nih.gov/sra/docs/sradownload/) command, and converted to fastq files using fastq-dump. Fastq quality control, data filtering and short read alignment was performed as above. Meta-analysis involved that read counts were normalised to equal read depth. Heatmap read density matrices and average read density plots were computed as described in the section ‘Read densities heatmaps and average plots’. Read density matrices were calculated for both strands separately when indicated. Ratio of XR-ser reads between directions and calculation of variability between directions was performed as described in legend of Fig. 6. ‘S-F’ scores and quantification of reads around TT loci was performed as in Lavigne et al^28^ with clusters borders defined previously.

### FANTOM5 Cap-analysis of Gene Expression (CAGE) sequencing data meta-analysis

The FANTOM5 strand specific CAGE-seq alignment files of normal Dermal fibroblast primary cells (6 Donors with source codes: 11269-116G9, 11346-117G5, 11418-118F5, 11450-119A1, 11454-119A5 and 11458-119A9) and normal skin fibroblasts (2 Donors with source codes: 11553-120C5 and 11561-120D4) were downloaded from ftp://ftp.biosciencedbc.jp/archive/fantom5/datafiles/phase2.2/basic/human.primary_cell.hCAGE and were combined. Heatmap read density matrices and average read density plots were computed as described in the section ‘Read densities heatmaps and average plots’. Read density matrices were calculated for both strands separately.

### Reactome pathway analysis

DAR-gain regions were associated with active transcripts, either directly by searching for genomic overlaps with annotated promoters (this resulted to 2,767 DAR-gain at promoter) or by searching genomic overlaps with intergenic/intragenic FANTOM5 enhancers (this resulted to 820 DAR-gain at enhancers). DAR-gain at enhancers were functionaly defined by analysing predicted enhancer-promoter associations retrieved from the FANTOM5 project site from ‘enhancer.binf.ku.dk/presets/human.associations.hdr.txt.gz’ and generated an additional 1,268 gene targets. A total of 3,284 of unique gene names were finally used to perform REACTOME pathway enrichment analysis using the R package ReactomePA: https://bioconductor.org/packages/release/bioc/html/ReactomePA.html.

## Data Availability

The data reported in this manuscript have been deposited with the Gene Expression Omnibus under accession code GSE125181, and will be released upon publication.

## Acknowledgements

We thank Pantelis Hatzis, Mihalis Verykokakis and members of the Fousteri lab for critical discussions and reading of the manuscript. We thank Vladimir Benes and the Genecore facility (EMBL, Germany) for the special care they use in sequencing our NGS libraries. This work was funded by a European Research Council grant to M.F., Agreement-309612 (TransArrest) and <Matching Funds> to MF funded by National sources.

## Author Contributions

M.F. and M.D.L. designed the study and were responsible for interpretation of the results. M.F. directed the study and obtained financial support. M.D.L., M.F., A.L. wrote the manuscript and all authors edited the manuscript. A.L. performed the experimental part of the study. D.K performed the statistical and bioinformatics analyses. M.D.L. contributed significantly in the bioinformatics analysis of the data. All authors discussed the results, reviewed, commented on and approved the final version of the manuscript. A.L and D.K. contributed equally to the paper.

## References

1. Haberle, V. & Stark, A. Eukaryotic core promoters and the functional basis of transcription initiation. Nature Reviews Molecular Cell Biology 19, 621–637 (2018).

2. Jonkers, I. & Lis, J. T. Getting up to speed with transcription elongation by RNA polymerase II. Nature Reviews Molecular Cell Biology 16, 167–177 (2015).

3. Lai, F. & Shiekhattar, R. Enhancer RNAs: the new molecules of transcription. Curr Opin Genet Dev 25, 38–42 (2014).

4. The FANTOM Consortium et al. An atlas of active enhancers across human cell types and tissues. Nature 507, 455–461 (2014).

5. Kim, T.-K. et al. Widespread transcription at neuronal activity-regulated enhancers. Nature 465, 182–187 (2010).

6. Preker, P. et al. RNA Exosome Depletion Reveals Transcription Upstream of Active Human Promoters. Science 322, 1851–1854 (2008).

7. Henriques, T. et al. Stable Pausing by RNA Polymerase II Provides an Opportunity to Target and Integrate Regulatory Signals. Molecular Cell 52, 517–528 (2013).

8. Compe, E. & Egly, J.-M. Nucleotide Excision Repair and Transcriptional Regulation: TFIIH and Beyond. Annual Review of Biochemistry 85, 265–290 (2016).

9. Lai, W. K. M. & Pugh, B. F. Genome-wide uniformity of human ‘open’ pre-initiation complexes. Genome Research 27, 15–26 (2017).

10. Nechaev, S. et al. Global Analysis of Short RNAs Reveals Widespread Promoter-Proximal Stalling and Arrest of Pol II in Drosophila. Science 327, 335–338 (2010).

11. Williams, L. H. et al. Pausing of RNA Polymerase II Regulates Mammalian Developmental Potential through Control of Signaling Networks. Molecular Cell 58, 311–322 (2015).

12. Cheng, B. & Price, D. H. Properties of RNA Polymerase II Elongation Complexes Before and After the P-TEFb-mediated Transition into Productive Elongation. Journal of Biological Chemistry 282, 21901–21912 (2007).

13. Core, L. & Adelman, K. Promoter-proximal pausing of RNA polymerase II: a nexus of gene regulation. Genes & Development (2019). doi:10.1101/gad.325142.119

14. Henriques, T. et al. Widespread transcriptional pausing and elongation control at enhancers. Genes & Development 32, 26–41 (2018).

15. Core, L. J. et al. Analysis of nascent RNA identifies a unified architecture of initiation regions at mammalian promoters and enhancers. Nature Genetics 46, 1311–1320 (2014).

16. Krebs, A. R. et al. Genome-wide Single-Molecule Footprinting Reveals High RNA Polymerase II Turnover at Paused Promoters. Molecular Cell 67, 411–422.e4 (2017).

17. Steurer, B. et al. Live-cell analysis of endogenous GFP-RPB1 uncovers rapid turnover of initiating and promoter-paused RNA Polymerase II. Proceedings of the National Academy of Sciences 201717920 (2018). doi:10.1073/pnas.1717920115

18. Jackson, S. P. & Bartek, J. The DNA-damage response in human biology and disease. Nature 461, 1071–1078 (2009).

19. Liakos, A., Lavigne, M. D. & Fousteri, M. Nucleotide Excision Repair: From Neurodegeneration to Cancer. in Personalised Medicine: Lessons from Neurodegeneration to Cancer (ed. El-Khamisy, S.) 17–39 (Springer International Publishing, 2017). doi:10.1007/978-3-319-60733-7_2

20. Helleday, T., Eshtad, S. & Nik-Zainal, S. Mechanisms underlying mutational signatures in human cancers. Nature Reviews Genetics 15, 585–598 (2014).

21. Hoeijmakers, J. H. J. DNA Damage, Aging, and Cancer. New England Journal of Medicine 361, 1475–1485 (2009).

22. Marteijn, J. A., Lans, H., Vermeulen, W. & Hoeijmakers, J. H. J. Understanding nucleotide excision repair and its roles in cancer and ageing. Nature Reviews Molecular Cell Biology 15, 465–481 (2014).

23. Vermeulen, W. & Fousteri, M. Mammalian Transcription-Coupled Excision Repair. Cold Spring Harbor Perspectives in Biology 5, a012625–a012625 (2013).

24. Hu, J., Adar, S., Selby, C. P., Lieb, J. D. & Sancar, A. Genome-wide analysis of human global and transcription-coupled excision repair of UV damage at single-nucleotide resolution. Genes & Development 29, 948–960 (2015).

25. Scharer, O. D. Nucleotide Excision Repair in Eukaryotes. Cold Spring Harbor Perspectives in Biology 5, a012609–a012609 (2013).

26. Cleaver, J. E., Lam, E. T. & Revet, I. Disorders of nucleotide excision repair: the genetic and molecular basis of heterogeneity. Nature Reviews Genetics 10, 756–768 (2009).

27. Djebali, S. et al. Landscape of transcription in human cells. Nature 489, 101–108 (2012).

28. Lavigne, M. D., Konstantopoulos, D., Ntakou-Zamplara, K. Z., Liakos, A. & Fousteri, M. Global unleashing of transcription elongation waves in response to genotoxic stress restricts somatic mutation rate. Nature Communications 8, 2076 (2017).

29. Williamson, L. et al. UV Irradiation Induces a Non-coding RNA that Functionally Opposes the Protein Encoded by the Same Gene. Cell 168, 843–855.e13 (2017).

30. Magnuson, B. et al. Identifying transcription start sites and active enhancer elements using BruUV-seq. Scientific Reports 5, 17978 (2015).

31. Donahue, B. A., Yin, S., Taylor, J. S., Reines, D. & Hanawalt, P. C. Transcript cleavage by RNA polymerase II arrested by a cyclobutane pyrimidine dimer in the DNA template. Proceedings of the National Academy of Sciences 91, 8502–8506 (1994).

32. Rockx, D. A. et al. UV-induced inhibition of transcription involves repression of transcription initiation and phosphorylation of RNA polymerase II. Proceedings of the National Academy of Sciences 97, 10503–10508 (2000).

33. Heine, G. F., Horwitz, A. A. & Parvin, J. D. Multiple Mechanisms Contribute to Inhibit Transcription in Response to DNA Damage. Journal of Biological Chemistry 283, 9555–9561 (2008).

34. Gregersen, L. H. & Svejstrup, J. Q. The Cellular Response to Transcription-Blocking DNA Damage. Trends in Biochemical Sciences 43, 327–341 (2018).

35. Geijer, M. E. & Marteijn, J. A. What happens at the lesion does not stay at the lesion: Transcription-coupled nucleotide excision repair and the effects of DNA damage on transcription in cis and trans. DNA Repair 71, 56–68 (2018).

36. Epanchintsev, A. et al. Cockayne’s Syndrome A and B Proteins Regulate Transcription Arrest after Genotoxic Stress by Promoting ATF3 Degradation. Molecular Cell 68, 1054–1066.e6 (2017).

37. Gyenis, Á. et al. UVB Induces a Genome-Wide Acting Negative Regulatory Mechanism That Operates at the Level of Transcription Initiation in Human Cells. PLoS Genetics 10, e1004483 (2014).

38. Andrade-Lima, L. C., Veloso, A., Paulsen, M. T., Menck, C. F. M. & Ljungman, M. DNA repair and recovery of RNA synthesis following exposure to ultraviolet light are delayed in long genes. Nucleic Acids Research 43, 2744–2756 (2015).

39. Bugai, A. et al. P-TEFb Activation by RBM7 Shapes a Pro-survival Transcriptional Response to Genotoxic Stress. Molecular Cell 74, 254–267.e10 (2019).

40. Chen, R. et al. PP2B and PP1 cooperatively disrupt 7SK snRNP to release P-TEFb for transcription in response to Ca2+ signaling. Genes & Development 22, 1356–1368 (2008).

41. Borisova, M. E. et al. p38-MK2 signaling axis regulates RNA metabolism after UV-light-induced DNA damage. Nature Communications 9, 1017 (2018).

42. Chiou, Y.-Y., Hu, J., Sancar, A. & Selby, C. P. RNA polymerase II is released from the DNA template during transcription-coupled repair in mammalian cells. Journal of Biological Chemistry jbc.RA117.000971 (2017). doi:10.1074/jbc.RA117.000971

43. Corces, M. R. et al. An improved ATAC-seq protocol reduces background and enables interrogation of frozen tissues. Nature Methods 14, 959 (2017).

44. Hauer, M. H. & Gasser, S. M. Chromatin and nucleosome dynamics in DNA damage and repair. Genes Dev. 31, 2204–2221 (2017).

45. Polo, S. E. & Almouzni, G. Chromatin dynamics after DNA damage: the legacy of the Access-Repair-Restore model. DNA Repair (Amst) 36, 114–121 (2015).

46. Misteli, T. & Soutoglou, E. The emerging role of nuclear architecture in DNA repair and genome maintenance. Nature Reviews Molecular Cell Biology 10, 243–254 (2009).

47. Zentner, G. E. & Henikoff, S. High-resolution digital profiling of the epigenome. Nature Reviews Genetics 15, 814–827 (2014).

48. Creyghton, M. P. et al. Histone H3K27ac separates active from poised enhancers and predicts developmental state. PNAS 107, 21931–21936 (2010).

49. Di Croce, L. & Helin, K. Transcriptional regulation by Polycomb group proteins. Nature Structural & Molecular Biology 20, 1147–1155 (2013).

50. Cifuentes-Rojas, C., Hernandez, A. J., Sarma, K. & Lee, J. T. Regulatory Interactions between RNA and Polycomb Repressive Complex 2. Molecular Cell 55, 171–185 (2014).

51. Kaneko, S., Son, J., Bonasio, R., Shen, S. S. & Reinberg, D. Nascent RNA interaction keeps PRC2 activity poised and in check. Genes Dev. 28, 1983–1988 (2014).

52. Chen, Y. et al. Principles for RNA metabolism and alternative transcription initiation within closely spaced promoters. Nature Genetics 48, 984–994 (2016).

53. Adar, S., Hu, J., Lieb, J. D. & Sancar, A. Genome-wide kinetics of DNA excision repair in relation to chromatin state and mutagenesis. Proceedings of the National Academy of Sciences 113, E2124–E2133 (2016).

54. Bugai, A. et al. P-TEFb activation by RBM7 shapes a pro-survival transcriptional response to genotoxic stress. bioRxiv (2018). doi:10.1101/394239

55. Mueller, B. et al. Widespread changes in nucleosome accessibility without changes in nucleosome occupancy during a rapid transcriptional induction. Genes Dev. 31, 451–462 (2017).

56. Karlic, R., Chung, H.-R., Lasserre, J., Vlahoviček, K. & Vingron, M. Histone modification levels are predictive for gene expression. PNAS 107, 2926–2931 (2010).

57. Gray, L. T. et al. Layer-specific chromatin accessibility landscapes reveal regulatory networks in adult mouse visual cortex. eLife 6, (2017).

58. Ucar, D. et al. The chromatin accessibility signature of human immune aging stems from CD8+ T cells. Journal of Experimental Medicine jem.20170416 (2017). doi:10.1084/jem.20170416

59. Beltran, M. et al. The interaction of PRC2 with RNA or chromatin is mutually antagonistic. Genome Res. 26, 896–907 (2016).

60. Schick, S. et al. Dynamics of chromatin accessibility and epigenetic state in response to UV damage. Journal of Cell Science 128, 4380–4394 (2015).

61. Huang, R.-P. et al. Egr-1 inhibits apoptosis during the UV response: correlation of cell survival with Egr-1 phosphorylation. Cell Death & Differentiation 5, 96–106 (1998).

62. Rehemtulla, A., Hamilton, C. A., Chinnaiyan, A. M. & Dixit, V. M. Ultraviolet Radiation-induced Apoptosis Is Mediated by Activation of CD-95 (Fas/APO-1). J. Biol. Chem. 272, 25783–25786 (1997).

63. Caricchio, R., McPhie, L. & Cohen, P. L. Ultraviolet B Radiation-Induced Cell Death: Critical Role of Ultraviolet Dose in Inflammation and Lupus Autoantigen Redistribution. The Journal of Immunology 171, 5778–5786 (2003).

64. Core, L. J. et al. Defining the Status of RNA Polymerase at Promoters. Cell Reports 2, 1025–1035 (2012).

65. Ibrahim, M. M. et al. Determinants of promoter and enhancer transcription directionality in metazoans. Nature Communications 9, 4472 (2018).

66. Scruggs, B. S. et al. Bidirectional Transcription Arises from Two Distinct Hubs of Transcription Factor Binding and Active Chromatin. Molecular Cell 58, 1101–1112 (2015).

67. Blasius, M., Wagner, S. A., Choudhary, C., Bartek, J. & Jackson, S. P. A quantitative 14-3-3 interaction screen connects the nuclear exosome targeting complex to the DNA damage response. Genes & Development 28, 1977–1982 (2014).

68. Tiedje, C. et al. p38 MAPK /MK2-mediated phosphorylation of RBM7 regulates the human nuclear exosome targeting complex. RNA 21, 262–278 (2015).

69. Boeing, S. et al. Multiomic Analysis of the UV-Induced DNA Damage Response. Cell Reports (2016). doi:10.1016/j.celrep.2016.04.047

70. Ehrensberger, A. H., Kelly, G. P. & Svejstrup, J. Q. Mechanistic Interpretation of Promoter-Proximal Peaks and RNAPII Density Maps. Cell 154, 713–715 (2013).

71. Adebali, O., Chiou, Y.-Y., Hu, J., Sancar, A. & Selby, C. P. Genome-wide transcription-coupled repair in Escherichia coli is mediated by the Mfd translocase. PNAS 114, E2116–E2125 (2017).

72. Murray, S. C. & Mellor, J. Using both strands: The fundamental nature of antisense transcription. Bioarchitecture 6, 12–21 (2016).

73. Seila, A. C., Core, L. J., Lis, J. T. & Sharp, P. A. Divergent transcription: A new feature of active promoters. Cell Cycle 8, 2557–2564 (2009).

74. Haradhvala, N. J. et al. Mutational Strand Asymmetries in Cancer Genomes Reveal Mechanisms of DNA Damage and Repair. Cell 164, 538–549 (2016).

75. Perera, D. et al. Differential DNA repair underlies mutation hotspots at active promoters in cancer genomes. Nature 532, 259–263 (2016).

76. Sabarinathan, R., Mularoni, L., Deu-Pons, J., Gonzalez-Perez, A. & López-Bigas, N. Nucleotide excision repair is impaired by binding of transcription factors to DNA. Nature 532, 264–267 (2016).

77. Shao, W. & Zeitlinger, J. Paused RNA polymerase II inhibits new transcriptional initiation. Nature Genetics 49, 1045–1051 (2017).

78. Fitz, J., Neumann, T. & Pavri, R. Regulation of RNA polymerase II processivity by Spt5 is restricted to a narrow window during elongation. The EMBO Journal 37, e97965 (2018).

79. Gressel, S. et al. CDK9-dependent RNA polymerase II pausing controls transcription initiation. eLife 6, e29736 (2017).

80. Xu, S., Grullon, S., Ge, K. & Peng, W. Spatial clustering for identification of ChIP-enriched regions (SICER) to map regions of histone methylation patterns in embryonic stem cells. Methods in molecular biology (Clifton, N.J.) 1150, 97–111 (2014).

81. Zhang, Y. et al. Model-based Analysis of ChIP-Seq (MACS). Genome Biology 9, R137 (2008).

